# Human footprints in the gut: how anthropogenic environments reshape the microbiome of chacma baboons

**DOI:** 10.64898/2026.06.10.731264

**Authors:** Célia Lacomme, Aluwani Ramaru, Benjamin Rey, Franck Prugnolle, Laure Segurel, Virginie Rougeron

## Abstract

Anthropogenic pressures are increasingly reshaping wildlife habitats worldwide. These transformations reduce natural areas, but also create new ecological niches, food resources, and environmental stressors, with potential consequences for wildlife behavior, physiology, and morphology. These changes may affect the gut microbiome, a critical component of host health, yet such effects are often inconsistent across species, particularly in wild non-human primates, and remain poorly understood. Here, we investigated how the gut microbiome of chacma baboons (*Papio ursinus ursinus*), an ecologically flexible generalist, responds to an anthropization gradient. We analyzed 512 fecal samples collected from 33 wild troops across a broad range of anthropogenic environments in the Western Cape, South Africa. Using a multi-metric approach including the Human Footprint Index, land-use variables and dietary proxies derived from stable isotopes, we assessed gut microbial diversity and composition based on 16S rRNA gene (V4) sequencing. Human-altered environments characterized by high Human Footprint and built-up areas were associated with reduced microbial diversity, and compositional and functional shifts, including decline in fiber-degrading taxa and increase in bacteria associated with simple carbohydrate and dairy metabolism. In contrast, highly cultivated areas showed no diversity difference and distinct microbial assemblages, while dietary variation had weaker effects, primarily altering rare taxa. Our results demonstrate that different components of anthropogenic pressure exert contrasting effects on the baboon gut microbiome, reflecting multiple ecological pathways extending beyond diet alone. Microbiome shifts may have implications for host health, potentially increasing susceptibility to pathogens or inflammatory diseases, with consequences for wildlife populations.

## Introduction

Anthropogenic activities are primary drivers of current environmental changes [1], with land-use conversion, infrastructure expansion, and urbanization being among the most disruptive alterations of terrestrial ecosystems [2,3]. The rapid expansion of human-modified landscapes leads to habitat loss and fragmentation [4], increased pollution [5], shifts in food resources [6], and contact with novel species and pathogens. Together, these changes generate novel selective pressures on wild populations [7].

Species differ markedly in their ability to cope with such pressures. Those with narrow ecological niches (e.g. specific diet and habitat), limited phenotypic plasticity, restricted dispersal, or slow life-history strategies are particularly vulnerable, often experiencing population decline and, ultimately, extinction [8,9]. Conversely, some species benefit from these anthropogenic environments through their ability to exploit novel resources and adjust rapidly to new conditions [10]. Human-modified environments may also reduce predation or competition, facilitating population persistence or expansion [11,12]. Species thriving in anthropized environments are numerous, including the red fox (*Vulpes vulpes*), common raccoon (*Procyon lotor*), hooded crow (*Corvus cornix*), and eastern chipmunk (*Tamias striatus*) [13–15]. However, persistence often comes with profound phenotypic changes, including altered behaviour [16], morphology (e.g. shorter beak and wing length [17,18]), and physiology (e.g. increased cholesterol or glucocorticoid levels [19–21]), potentially leading to long-term fitness costs [22]. Thus, understanding how species respond to anthropogenic environments is critical for predicting biodiversity changes.

One of the key host-associated systems expected to mediate these responses is microbial communities. Microorganisms indeed play fundamental roles in ecosystem functioning and animal health [23]. In particular, the gut microbiota – the microorganisms inhabiting the gastrointestinal tract - play a central role in host physiology by influencing host metabolism [24,25], digestion [26], immune function [27], brain development [28], and behaviour [29]. Beyond its effects on host physiology, the gut microbiota is itself shaped by a combination of intrinsic host characteristics (e.g. genetics, age, sex, immune status) and extrinsic drivers (including diet, seasonality, and habitat quality) [30,31]. Importantly, the gut microbiota is highly plastic, responding to environmental change through shifts in diversity and/or composition. It is therefore increasingly recognized as a key mediator of short- and long-term host adaption [32,33]. This interplay has created growing interest in considering the gut microbiome in wildlife health and conservation in human-modified environments [30,34].

Despite increasing research on how anthropization impacts wildlife gut microbiome, previous studies have revealed inconsistent patterns across host species, reflecting important knowledge gaps. Most studies report a decrease in microbial diversity in human-altered environments, such as passerines in urban areas [35], lemurs in disturbed forests [36], or bats in monocultures [37]. In contrast, other studies reported increased diversity, including in coyotes in urban areas [38], and black howler monkeys inhabiting disturbed forests [39], while no detectable effect was observed in chameleons in urban environments [40]. In addition, anthropization can also induce shifts in microbial composition, particularly in primates. Studies on macaques, red colobus, and brown howler monkeys have reported compositional restructuring associated with habitat disturbance or human provisioning [41– 43]. These results likely highlightx the ecological complexity and methodological differences across microbiome studies. The divergent responses may arise from differences in anthropogenic disturbance definition, often relying on categorical classifications (urban vs natural) and lacking quantitative measures [44,45]. Furthermore, inconsistencies among studies may also emerge from limited replication across contrasting habitats, often relying on a single population per habitat. Importantly, anthropogenic environments are highly heterogeneous, encompassing multiple landscape types and intensities of human pressures beyond urbanization alone, which may differentially shape microbial communities. Together, these limitations highlight the need for well-replicated studies across multiple populations, combined with quantitative metrics of anthropogenic pressure, in systems that allow to robustly assess how human-modified environments shape gut microbial communities.

Chacma baboons (*Papio ursinus*) represent a relevant model for investigating anthropogenic impacts, as they inhabit diverse environments [46] and are among the most generalist non-human primates, with an omnivorous diet [47]. This flexibility enables them to colonize human-modified habitats (e.g. disturbed forests, agricultural land, peri-urban and urban areas) and consume human-derived foods, as is the case in South Africa [48,49], frequently leading to human–baboon conflicts [50]. These environments may affect their bacterial communities directly through exposure to novel microbes, or indirectly via changes in diet, behaviour, stress, or immune function [51]. Indeed, the baboon gut microbiome is driven by environmental factors such as diet, seasonality, and soil properties (yellow baboon [52,53]; olive baboon [54]). Anthropogenic pressures, specifically habitat fragmentation, have been associated with reduced microbial diversity and shifts in composition in yellow baboon [42,55], and captivity in chacma baboon led to a humanization of their gut microbiome [56]. Despite these insights, studies have largely focused on isolated populations or specific environmental factors, lacking the quantitative resolution needed to disentangle the effects of varying degrees and types of anthropization. Studying baboon populations along a continuum of anthropogenic environments provides a powerful framework for investigating how human-modified environments shape gut microbial communities.

In this study, we investigated how the gut microbiome of Cape chacma baboons (*Papio ursinus ursinus*) varies along a continuous anthropogenic gradient in the Western Cape, South Africa. We analysed 512 fecal samples from 33 wild troops across contrasted environments ranging from protected parks to agricultural land, peri-urban, and urban areas. To our knowledge, this is the first large-scale study of baboon gut microbiome encompassing such a diversity of anthropogenic contexts. To move beyond discrete categories, we quantified anthropization using multiple parameters: a broad-scale combined measure of human pressure, the Human Footprint [57], and finer-scale land-use indicators and stable isotope values, reflecting baboon habitat and diet, respectively. This multi-metric approach aimed to evaluate the overall effect of anthropization on chacma baboons’ microbial community diversity and composition, and the respective contributions of land-use and diet, with three predictions. We expected (1) reduced gut microbial diversity with increasing anthropogenic pressure, with stronger effects in individuals experiencing pronounced dietary changes; (2) shifts in community composition along the anthropogenic gradient, with more homogeneous microbial communities in less anthropized environments and higher variability in highly modified habitats; and (3) taxonomic and functional changes consistent with dietary transitions, including reduced abundance or loss of fiber-associated taxa and an enrichment of taxa linked to fat energy-dense or human-associated diets [41–43,54,55]. Overall, this study provides key insights into how different types and intensities of anthropization shape gut microbial communities in chacma baboons, and, more broadly, primate responses to human-modified environments.

## Materials and methods

### Ethics

This project was approved by the Nelson Mandela University Research Ethics Committee (Permit no. 0544), and by the relevant local authorities, including Cape Nature (CN44-87-28483), SANparks (CRC/2024-2025/008), municipalities (George, Knysna, Bitou, Overstrand and Cape Town), and under Section 20 of the Animal Diseases Act, 1984 (12/11/1/6/1; [6285 CM]).

### Study sites and troops

This study was conducted on 33 wild chacma baboon troops along the southern coast of the Western Cape, South Africa. Nineteen troops were sampled in the Garden Route area (referred as GAR; George to Nature’s Valley) and 14 in the Cape Peninsula–Hermanus area (referred as CAPE) (Fig. 1A). These areas are characterized by a Mediterranean climate, with mainly forested habitats in GAR and shrubland in CAPE (Fig. 1B-C; Supplementary Table 1). Chacma baboons in this region inhabit a gradient of habitats, from relatively undisturbed environments to anthropogenically modified landscapes (rural, agricultural and urban areas), exploiting both natural and anthropogenic food resources.

**Figure 1.**
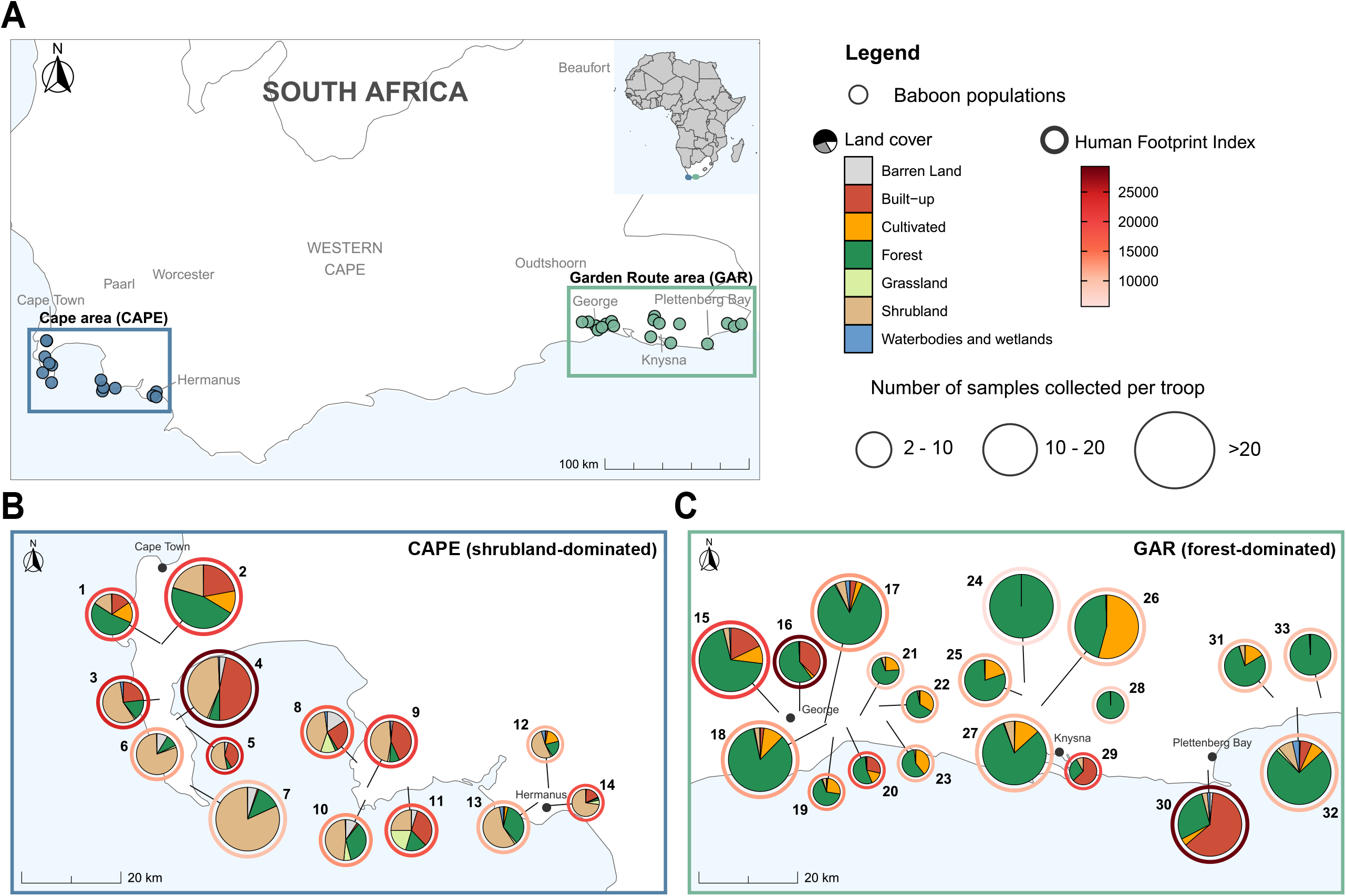
Study area and landscape characterization of chacma baboon troop habitats along an anthropization gradient in the Western Cape, South Africa. A. Location of the study areas within South Africa. Each point represents the baboon troop sampled in the Cape (n = 14) and GAR (n = 19) areas. B. Detailed map of the Cape Peninsula–Hermanus area (CAPE). C. Detailed map of the Garden Route area (GAR). Each pie chart illustrates the land cover composition within a 2km^2^ buffer for each baboon troop, based on the South African National Land Cover dataset (SANLC, 2022 – EPSG4326). The outer ring surrounding each pie chart represents the mean Human Footprint Index [59] in each buffer, reflecting cumulative anthropogenic pressure. The size of each pie chart is proportional to the number of fecal samples collected for troop. Numbers indicate baboon troop identities as listed in Supplementary Table 1.

### Sample collection

A total of 512 fecal samples were collected non-invasively from 33 troops of chacma baboons between March and May 2024. Each sample was split into two subsamples: one preserved in RNAlater (Thermo Fisher Scientific) for microbiome analysis and one without preservative for stable isotope analysis. For each sample, GPS location, date, time, and Bristol Stool Scale [58] were recorded. To control for potential environmental contamination, 52 soil samples were collected near fecal samples across substrate types. All samples were kept at 4°C during fieldwork and transferred to -20°C within 12h of collection at Nelson Mandela University Molecular laboratory until further processing.

### Anthropization characterization

Anthropogenic pressures were characterized using a two-step framework combining broad- and fine-scale metrics. First, we used the Human Footprint Index (HFI) as a proxy for overall human disturbance, integrating land cover change, human population density, electrical infrastructure, road and railway networks, and navigable waterways, with values ranging from 0 (no human pressure) to 50 (maximum pressure). For each troop, HFI was calculated as the mean value within a 1km radius buffer around sampling locations, using QGIS 3.38.3 and the 2020 Human Footprint dataset [59]. Buffers were aggregated at the troop level and reflect baboon minimal daily ranging distance [60].

Second, to obtain a finer-scale characterization of anthropogenic pressures, we used land-use variables and stable isotope values (δ^1^□N and δ^13^C), reflecting habitat and diet, respectively. Land-use variables included the proportions of eight land cover categories (built-up, barren land, cultivated land, forest, grassland, shrubland, waterbodies and wetlands) based on the 2022 South African National Land Cover Dataset [61], as well as road length and building density within each buffer, derived from QGIS and RStudio. Road and building data were obtained from the Global Mapping dataset and the Google Open Buildings, respectively [62]. Stable isotope analysis of nitrogen (δ^1^□N) and carbon (δ^13^C) from fecal samples reflect short-term dietary intake [63]. δ^1^□N provides a proxy for protein consumption, while δ^13^C discriminates among plant functional types (C3, C4, CAM) [63]. Samples were prepared following standard protocols [64] (Supplementary Materials and Methods).

### DNA extraction, 16S sequencing and sex characterization

DNA was extracted from 512 fecal samples and 52 soil samples using the Qiagen DNeasy PowerSoil Pro kit (QIAGEN, Germany), following the manufacturer’s protocol. The V4 region of the 16S rRNA gene was amplified using the primers 515F/806R with a dual-indexing approach [65] and the amplicons were sequenced on an Illumina MiSeq platform (V3, 2 × 300 bp paired-end), in two independent runs (Supplementary Materials and Methods). Extraction blanks (39), PCR negative controls (39), and sequencing blanks (9) were included to monitor contamination. To assess batch effects, ten samples were sequenced in duplicate across runs. Individual sex was determined using a multiplex PCR targeting the UTX, UTY and SRY regions, following the protocol described in [66] (Supplementary Materials and Methods).

### Bioinformatic processing

Eight samples failing sequencing were removed, leaving 504 fecal samples for analyses (plus 10 sequencing replicates). Sequences were processed in QIIME2 (v.2024.10) [67], using Cutadapt for primer removal and DADA2 for denoising (trimming: 240/160 bp; Q>30) and generating an amplicon sequence variant (ASV) table [68]. Taxonomy was assigned using the GreenGenes2 database (v.2024.09) [69] and mitochondrial and chloroplast sequences were excluded, resulting in 6,417 ASVs. A total of 23,801,111 reads were retained (mean: 46,396 per sample; range: 14,334 – 73,034) (Supplementary Materials and Methods). Phylogenetic trees were constructed using MAFFT and FastTree within the QIIME2 pipeline [67].

Data filtration included the exclusion of negative controls with negligible read counts (< 3 reads/sample), removal of 57 soil-associated ASVs (identified by higher relative abundance in soil than in feces), and exclusion of five non-baboon samples confirmed by mitochondrial 16S rRNA Sanger sequencing (Supplementary Fig. 1-2, Materials and Methods). Sequencing replicates and duplicate samples were removed based on high similarity (Bray-Curtis < 0.25), retaining only the sample with the highest read count (Supplementary Table 2).

Finally, ASVs with fewer than 50 total reads across samples were removed to eliminate very rare sequences. Data were rarefied to 25,000 reads per sample, yielding a final dataset of 438 fecal samples and 3,551 ASVs, with an average of 469 ASVs (± 78 SD) per sample. Summary statistics of sample and sequence retention across filtering steps are provided in the Supplementary Fig. 3.

### Statistical analyses

Analyses were performed using R [70] and figures were generated using ggplot2 [71] and edited in Inkscape. Additional methodology is provided in the Supplementary Materials and Methods.

#### Alpha diversity

Alpha diversity was assessed using observed richness, Shannon diversity, and Faith’s phylogenetic diversity (Faith’s PD), calculated with phyloseq and vegan packages [72,73]. Effects of anthropization were tested using linear and generalized linear mixed models (LMMs/GLMMs; lme4 [74]), with troop as a random effect. Two model structures were tested, including either HFI or land-use proportions and stable isotopes as predictors. δ^1^□N was decomposed into between- and within-troop components (δ^1^□N_mean and δ^1^□N_diff) to account for contrasting effects (Supplementary Fig. 4-5), while δ^13^C was included as a single variable. Sex, Bristol Stool Scale, and area were included as covariates. Collinearity was assessed using variance inflation factors (VIF), retaining built-up and cultivated variables (Supplementary Fig. 6). Model selection relied on model averaging (ΔAICc < 2). Model assumptions and spatial autocorrelation were assessed using DHARMa [75].

#### Beta diversity

Inter-individual variation in gut microbial community composition was assessed using weighted and unweighted UniFrac distances (phyloseq [72]). Effects of anthropization were tested using PERMANOVA (*adonis2*, vegan [73]) under the same two modelling frameworks as for alpha diversity. To account for non-independence among samples, permutations were restricted to the troop level (9,999 permutations). Spatial structure was assessed using Mantel tests between UniFrac dissimilarities and geographical distance matrices. Distance-based redundancy analysis (dbRDA [73]) was used to evaluate associations between microbial composition and predictor variables, controlling for spatial autocorrelation using dbMEMs [76]. Beta diversity dispersion was assessed using PERMDISP2 (betadisper, vegan [73]). Distances to centroids were calculated at the troop level (only for troops with more than 10 samples), and modelled using GLMMs (lme4 [74]), with HFI, sex, and area as fixed effects and troop as a random effect.

#### Differential abundance analysis

Differential taxa abundance across the anthropogenic gradient was assessed using ANCOM-BC2 [77] on the unrarefied dataset under both modelling frameworks. Only ASVs detected in at least 10% of samples were retained. P-values were corrected using the Benjamini–Hochberg procedure [78], and ASVs with adjusted p-values < 0.05 were considered differentially abundant. Analyses were repeated at the genus, family, and phylum levels.

#### Microbial functional prediction

Microbial functional profiles were predicted from 16S rRNA data (unrarefied dataset) using PICRUSt2 with default parameters [79], resulting in Enzyme Commissions (EC) and MetaCyc pathways. Differential abundance of functional pathways across the anthropization gradient was assessed under both modelling frameworks.

## Results

### Baboon gut microbiome characteristics

The majority of ASVs of the gut microbiome of chacma baboons belonged to Bacteria, with only eight reads (0.6%) assigned to Archaea. At the phylum level, the microbiome was dominated by two bacterial phyla: Bacillota_A (51.3%) and Bacteroidota (16.3%), followed by Pseudomonadota (6%), Spirochaetota (5%), Verrucomicrobiota (5%) (Supplementary Fig. 7A). At the family level, the community was primarily composed of Lachnospiraceae (15%), Bacteroidaceae (12%), Oscillospiraceae (11%), and Ruminococcaceae (8%) (Supplementary Fig. 7B). At the genus level, Prevotella was the most abundant taxon overall (11%), although relative abundances varied markedly across troops (Supplementary Fig. 7C).

### Effects of area, sex, and stool consistency on gut microbiome

All three alpha diversity metrics differed significantly between areas, with higher diversity in GAR compared to CAPE (observed richness, β = 0.09, 95% CI [0.03; 0.16]; Shannon diversity, β=0.11, 95% CI [0.01; 0.20]; Faith’s PD, β =1.36, 95% CI [0.22; 2.49]) (Table 1A, Supplementary Fig. 8A-C). Gut microbial composition also differed between areas (unweighted-UniFrac, R^2^ = 0.047, p < 0.001; weighted-UniFrac, R^2^ = 0.044, p < 0.001) (Supplementary Table 3A and Fig. 8D-E), revealing notably higher relative abundances of *Pararoseburia* and *Escherichia* and lower abundances of *Catenibacterium* in GAR compared to CAPE. Sex was also associated with differences in microbiome (Table 1A, Supplementary Fig. 9A-C), with males showing lower microbial diversity than females across all metrics (observed richness, β = -0.06, 95% CI [-0.09; -0.03]; Shannon diversity, β = -0.12, 95% CI [-0.18; -0.06]; Faith’s PD, β = -1.11, 95% CI [-1.66; -0.57]) (Table 1A). Smaller differences in composition were also observed (unweighted UniFrac, R^2^ = 0.007, p < 0.001; weighted-UniFrac, R^2^ = 0.005, p < 0.001) (Supplementary Table 3A and Fig. 9D-E), mainly driven by increases of *Ruminococcus, Anaerovibrio*, and *Faecalibacterium* in males.

**Table 1.** Linear mixed model results predicting variation in gut microbial alpha diversity in chacma baboons (n = 438 samples from 33 troops). (A) Models assessing the effect of the Human Footprint Index, and (B) models including land-use variables (built and cultivated areas) and stable isotopes (δ^1^□N and δ^13^C). The table reports model-averaged estimates and confidence intervals across models with ΔAICc < 2.

| A |  |  |  | B |  |  |  |
| --- | --- | --- | --- | --- | --- | --- | --- |
| Metrics | Predictors | Estimates | 95% confidence intervals | Metrics | Predictors | Estimates | 95% confidence intervals |
| Richness | Human Footprint | -0.04 | [-0.07; -0.01] | Richness | Built-up | -0.01 | [-0.05; 0.02] |
|  | Area-GAR | 0.09 | [0.03; 0.16] |  | Cultivated | 0.03 | [-0.05; 0.07] |
|  | Sex-Male | -0.06 | [-0.09; -0.03] |  | d15N_mean | -0.02 | [-0.05; 0.008] |
|  | Bristol scale | -0.02 | [-0.04; -0.01] |  | d15N_diff | 0.03 | [0.01; 0.04] |
| Shannon diversity | Human Footprint | -0.11 | [-0.16; -0.06] |  | d13C | -0.01 | [-0.02; 0.01] |
|  | Area-GAR | 0.11 | [0.01; 0.20] |  | Area-GAR | 0.10 | [0.03; 0.17] |
|  | Sex-Male | -0.12 | [-0.18; -0.06] |  | Sex-Male | -0.06 | [-0.09; -0.03] |
|  | Bristol scale | -0.03 | [-0.07; -0.01] |  | Bristol scale | -0.03 | [-0.04; -0.01] |
| Faith's phylogenetic diversity | Human Footprint | -0.65 | [-1.37; 0.08] | Shannon diversity | Built-up | -0.06 | [-0.11; -0.001] |
|  | Area-GAR | 1.36 | [0.22; 2.49] |  | Cultivated | 0.03 | [-0.03; 0.10] |
|  | Sex-Male | -1.11 | [-1.66; -0.57] |  | d15N_mean | -0.04 | [-0.09; 0.01] |
|  | Bristol scale | -0.4 | [-0.69; -0.12] |  | d15N_diff | 0.05 | [0.02; 0.07] |
|  |  |  |  |  | d13C | -0.02 | [-0.06; 0.01] |
|  |  |  |  |  | Area-GAR | 0.13 | [0.02; 0.25] |
|  |  |  |  |  | Sex-Male | -0.11 | [-0.17; -0.05] |
|  |  |  |  |  | Bristol scale | -0.05 | [-0.08; -0.02] |
|  |  |  |  | Faith's phylogenetic diversity | Built-up | -0.18 | [-0.81; 0.45] |
|  |  |  |  |  | Cultivated | 0.50 | [-0.18; 1.19] |
|  |  |  |  |  | d15N_mean | -0.24 | [-0.80; 0.31] |
|  |  |  |  |  | d15N_diff | 0.49 | [0.25; 0.73] |
|  |  |  |  |  | d13C | -0.23 | [-0.5; 0.07] |
|  |  |  |  |  | Area-GAR | 1.63 | [0.37; 2.88] |
|  |  |  |  |  | Sex-Male | -1.03 | [-1.56; -0.49] |
|  |  |  |  |  | Bristol scale | -0.48 | [-0.76; -0.19] |

Finally, stool consistency was negatively correlated with all alpha diversity metrics (Table 1A), with softer stools associated with lower diversity.

### Anthropogenic impact on gut microbiota alpha diversity

HFI had a significant negative effect on all alpha diversity metrics (Fig. 2A-C, Table 1A), with diversity declining with increasing Human Footprint (observed richness, β = –0.04, 95% CI = [-0.07; -0.01]; Shannon diversity, β = -0.11, 95% CI = [-0.16; -0.06]; Faith’s PD, β = -0.65, 95% CI = [-1.37; 0.08]). Model selection supported HFI as a predictor, with no interaction with sex or area retained.

**Figure 2.**
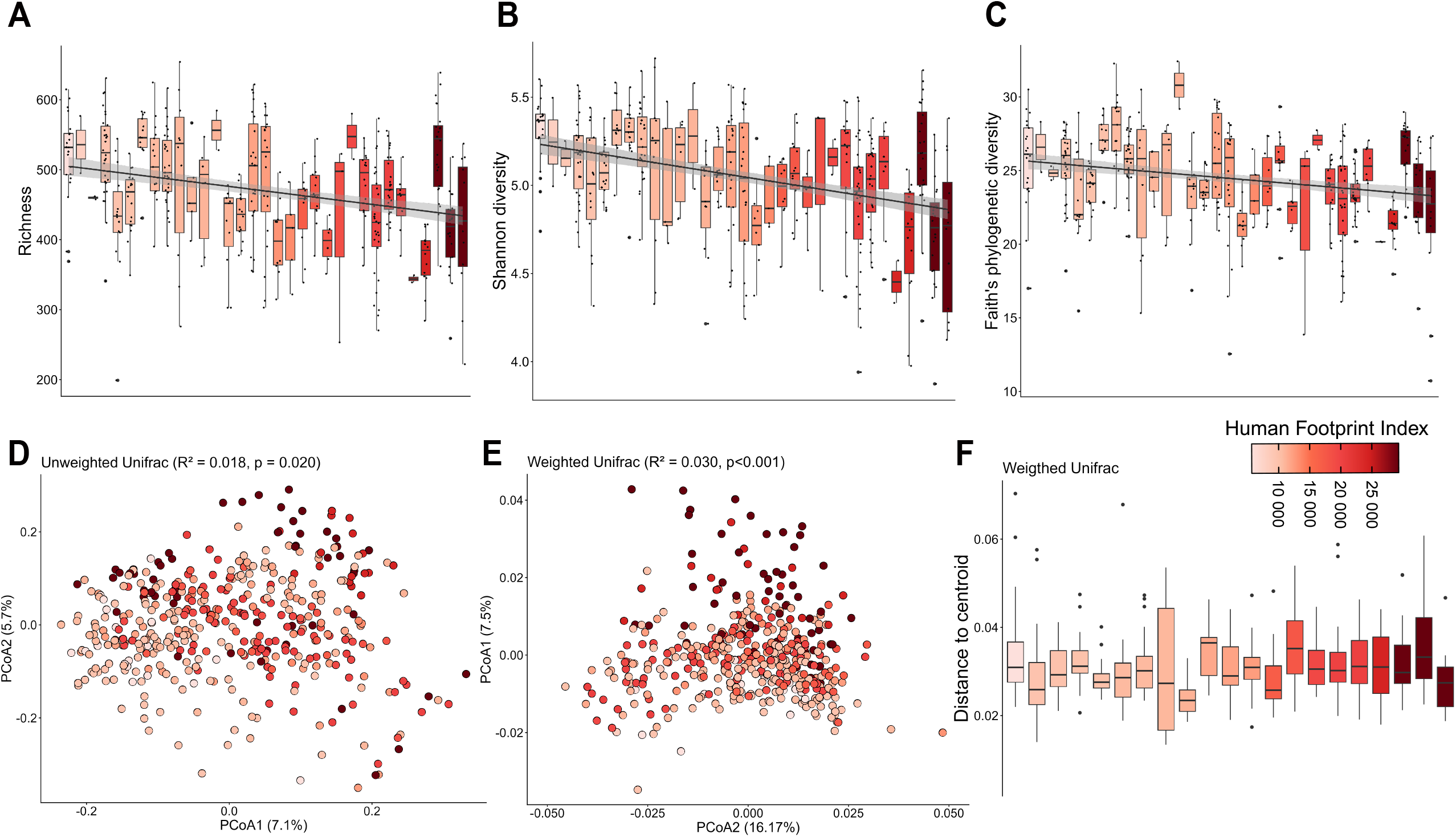
Effect of the Human Footprint Index on the diversity and composition of the baboon gut microbiome. Alpha diversity metrics are shown at the troop level for (A) richness, (B) Shannon diversity, and (C) Faith’s phylogenetic diversity. Solid lines represent linear regressions with shaded areas indicating the standard errors. Beta diversity patterns are represented using PCoA based on (D) unweighted UniFrac, and (E) weighted UniFrac distances. (F) Within-troop microbiome dispersion based on weighted UniFrac distances, measured as the distance to centroid (no significant association with HFI). All panels are coloured according to the Human Footprint Index. For panels A–E, N = 438 samples across 33 troops. Panel F includes 21 troops after excluding troops with fewer than 10 samples.

In the fine-scale framework, built-up proportion was negatively associated with Shannon diversity, with no significant effects on ASV richness or Faith’s PD (Table 1B; β = -0.06, 95% CI = [-0.11; -0.001]). Cultivated areas had no significant effect on alpha diversity (Table 1B). For the diet-related predictors, mean troop-level δ^1^□N and δ^13^C values, although retained in the best-supported models, were not significantly associated with alpha diversity (Table 1B). In contrast, within-troop δ^1^□N variation (δ^1^□N_diff) was positively associated with all three metrics (Table 1B). No residual spatial autocorrelation was detected in any model.

### Gut microbiota composition changes along an anthropogenic gradient

HFI significantly influenced gut microbial composition for both unweighted and weighted UniFrac dissimilarities (unweighted UniFrac, R^2^ = 0.018, p = 0.020; weighted UniFrac, R^2^ = 0.030, p<0.001; Fig. 2D-E, Supplementary Table 3A). Similarly, built-up and cultivated variables were significantly associated with microbial composition changes across both metrics (Supplementary Table 3B). Among dietary predictors, δ^1^□N (between troops variation) significantly influenced the microbial composition for unweighted UniFrac (R^2^ = 0.018, p = 0.005) and marginally for weighted UniFrac (R^2^ = 0.016, p = 0.053), and δ^13^C (between troops variation) was only significantly associated with unweighted UniFrac (R^2^ = 0.011, p = 0.032) (Supplementary Table 3B). No significant interaction between sex and the environmental variables was detected.

As beta diversity showed significant spatial structure (Mantel tests, p < 0.01), we evaluated whether these associations remained significant after controlling for spatial autocorrelation. HFI remained significant for weighted (p = 0.011), but not for unweighted Unifrac (Supplementary Table 4A). Built-up areas and δ^1^□N showed marginal effects on weighted UniFrac, p = 0.071 and 0.072), while δ^1^□N and δ^13^C were still significant for unweighted UniFrac (p = 0.039 and 0.015). All other tests were not significant (Supplementary Table 4B).

Finally, HFI had no significant effect on community dispersion, regardless of the UniFrac metric (Fig. 2F, Supplementary Table 5).

### Differential abundances of microbial taxa along an anthropogenic gradient

We then identified specific bacterial taxa contributing to the observed patterns. Overall, 8/24 phyla, 25/153 families, 46/419 genera and 174/3,010 ASVs were significantly associated with HFI, whereas built-up areas showed fewer associated taxa (4 phyla, 15 families, 35 genera, and 121 ASVs) (Supplementary Table 6A-H). Consistent with the beta diversity results, HFI and built-up area showed concordant patterns, sharing 19 genera and 64 ASVs that shifted in the same direction (Fig. 3A, Supplementary Fig. 10). Increasing HFI and built-up proportions were consistently associated with higher abundances of Bifidobacteriaceae (*Bifidobacterium*) and Succinivibrionaceae (*Succinivibrio*), followed by Treponemataceae (*Treponema*) and Lactobacillaceae *(Ligilactobacillus)*. In contrast, several taxa declined, including *Faecalibacillus*, Acidaminococcaceae *(Phascolarctobacterium)*, and Lachnospiraceae (*Eubacterium)*. Some responses were specific to HFI, including an enrichment of Acidaminococcus and depletion of Lachnospiraceae (*Blautia, Coprococcus, Lachnoclostridium*), and Ruminococcaceae *(Gemmiger, Pygmaiobacter)*.

**Figure 3.**
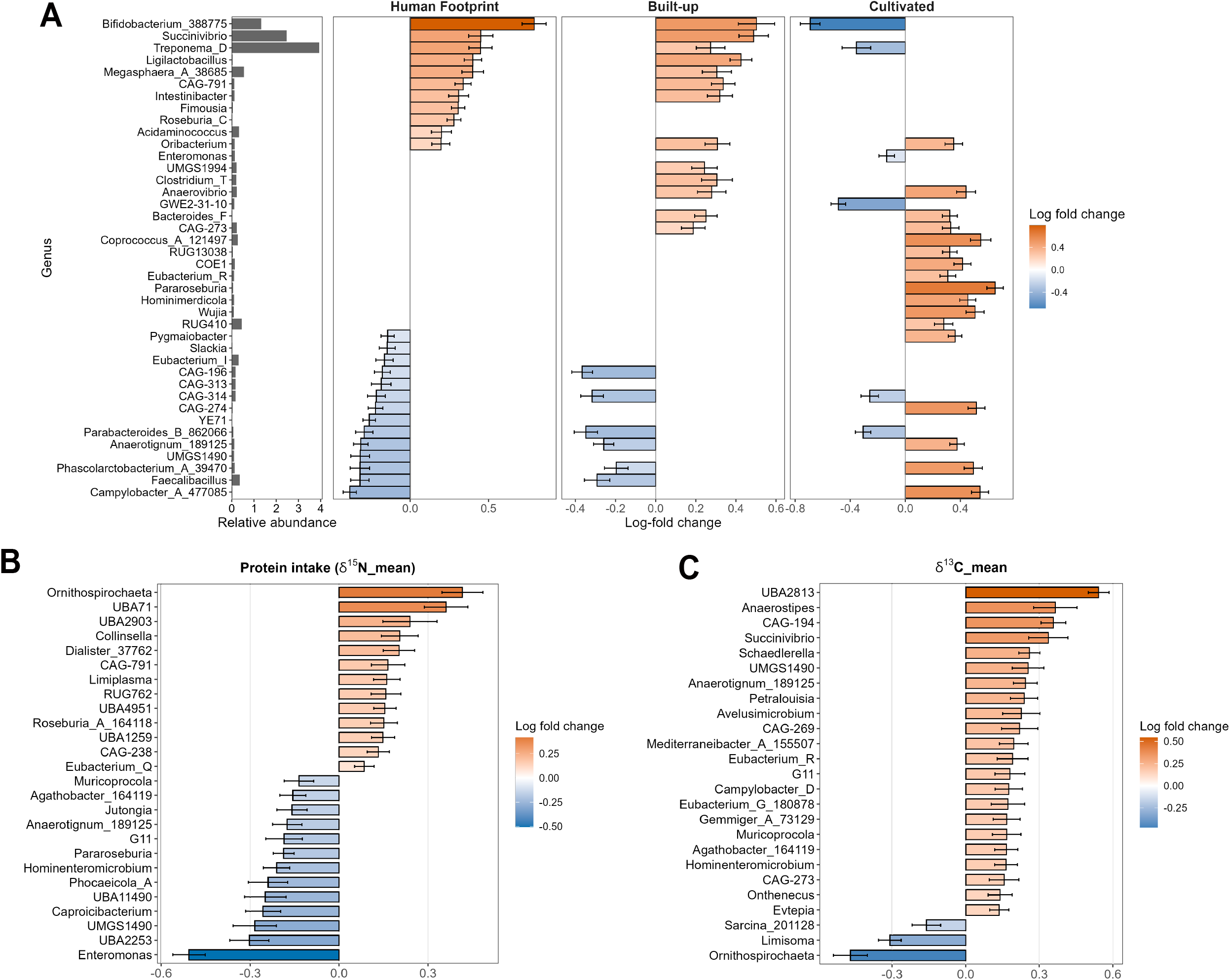
Differential abundance of microbial genera in chacma baboon along different anthropogenic variables. (A) The 40 genera showing the strongest differential abundance (log fold-change) in relation to Human Footprint, built-up and cultivated areas. The panel on the left shows the relative abundance of each genus. (B) Genera significantly associated with δ^1^□N values. (C) Genera significantly associated with δ^13^C values. The colour gradient corresponds to the magnitude of the differences (log fold-change), with increases shown in red and decreases in blue for each predictor. White cells indicate non-significant associations.

In contrast, cultivated areas exhibited distinct microbial signatures (4 phyla, 15 families, 49 genera and 160 ASVs). Unlike HFI and built-up, cultivated landscapes were associated with higher abundances of *Coprococcus, Campylobacter, Pygmaiobacter*, and *Phascolarctobacterium* and lower abundances of *Bifidobacterium* and *Treponema*, largely driven by the same ASVs (Fig. 3A). Despite these differences, *Bacteroides* and *Anaerovibrio* showed consistent positive associations with both built-up and cultivated land-use, whereas *Eubacterium_G* showed a consistent negative association. Additional shifts in cultivated land involved low-abundance taxa unaffected by HFI, including increases in Lachnospiraceae (*Butyrivibrio, Pararoseburia*), Acidaminococcaceae (*Hominimerdicola*), and Butyricicoccaceae (*Butyricicoccus*), and decreases in Enterobacteriaceae (*Escherichia*).

Dietary predictors (δ^1^□N and δ^13^C) affected fewer taxa (2 phyla, 8 families, 26 genera, and 114 ASVs, and 2 phyla, 5 families, 25 genera, and 78 ASVs, respectively, Fig. 3B-C). Interestingly, δ^1^□N showed four concordant genera overlapping with HFI (e.g. increase in *Muricoprocola*), while δ^13^C shared five concordant genera overlapping with cultivated areas (e.g. increase in *Agathobacter*) (Supplementary Fig. 10). Additionally, higher δ^1^□N was associated with significantly higher abundances of *Collinsella, Dialister, and Roseburia*, and lower abundances of *Enteromonas, Caproicibacterium*, and *Muricoprocola*, while higher δ^13^C was predominantly associated with enrichment of several taxa, including *Anaerostipes, Succinivibrio, Campylobacter, and Eubacterium*.

*Parabacteroides* was the only genus negatively associated with all land-use variables and HFI. Moreover, *Anaerotignum* (Clostridiaceae) was the only genus responding across all variables, decreasing with HFI, built-up areas and δ^1^□N, and increasing with cultivated land and δ^13^C.

### Metabolic functions associated with anthropogenic levels in chacma baboons

Predicted functional profiles of the microbial community were significantly associated with HFI, with 109/497 metabolic pathways and 647/2,589 enzyme functions showing differential abundances (Supplementary Table 6I-K).

High HFI showed enrichment of central carbohydrate metabolism pathways, including glycolysis, Entner–Doudoroff pathway, and glycogen degradation, as well as reductive tricarboxylic acid cycle and osmoprotectant utilization (e.g. ectoine and glycine betaine degradation) (Fig. 4). In contrast, several pathways showed reduced relative abundance with increasing HFI, including those involved in degradation of environmental pollutants (glyphosate, toluene, catechol, and salicylate), recycling of complex organic substrates (e.g. chitin deacetylation, fatty acid β-oxidation, and allantoin degradation), plant-derived biomass degradation (D-xylose), osmotic stress responses (ectoine biosynthesis) and amino acid metabolism, particularly the degradation of L-histidine and L-glutamate (Fig. 4).

**Figure 4.**
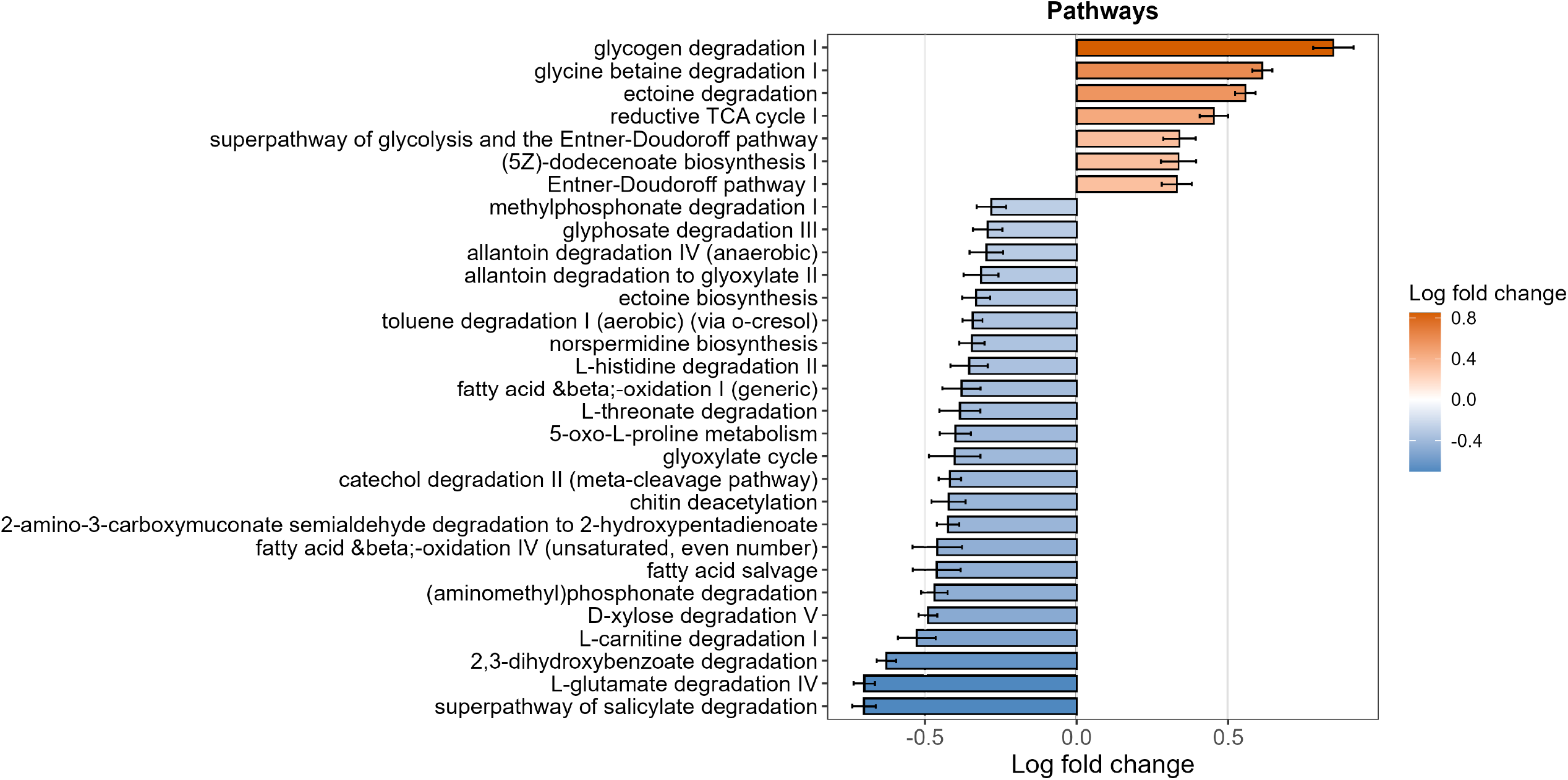
Differential abundance of metabolic pathways in chacma baboons along the Human Footprint Index. Values represent log fold-change, with increases shown in red and decreases in blue. The 30 pathways with the largest absolute LFC values are shown.

## Discussion

In this study, we investigated how gut microbial communities in chacma baboons vary along a continuous gradient of anthropogenic environments. Our multi-metric approach revealed that microbial responses differ depending on the type and level of anthropogenic disturbance. First, anthropization, captured by HFI and built-up areas, was associated with reduced gut microbial diversity, and marked compositional and functional shifts. Specifically, baboons in anthropogenic environments exhibited lower relative abundance of fiber-degrading taxa and higher abundance of taxa involved in the metabolism of carbohydrates, glycogen and dairy products. Second, baboons inhabiting cultivated areas showed distinct microbial signatures, suggesting ecological pressures differing from those in urban environments. Finally, dietary variations inferred from stable isotopes had a weaker and more targeted effect, primarily altering the presence of rare taxa.

### Baboon characteristics and environmental drivers of gut microbiome variation

Differences in microbial diversity and composition between the forested GAR area and fynbos-dominated CAPE area likely reflect habitat-specific ecological drivers. While forest ecosystems may offer richer microbial reservoirs due to higher moisture and biodiversity [80,81], these patterns likely reflect broader environmental differences, highlighting the importance of habitat context in microbiome studies. Gut microbial communities differed between males and females in both diversity and composition, with females exhibiting higher diversity, consistent with previous findings in primates and humans [55,82]. These differences may arise from differences in physiology (e.g. hormones), nutritional requirements (particularly during pregnancy and lactation), body mass, and behaviours such as foraging strategies or social interactions [82]. Dietary differences are unlikely to explain these sex differences, as δ^1^□N and δ^13^C did not differ between sexes, although variations in macronutrient composition or food processing not captured by these proxies cannot be excluded. Further studies are needed to explicitly test these mechanisms and clarify the drivers of sex-related variation in the gut microbiome. Stool consistency was significantly associated with microbial alpha diversity, reflecting gut transit time and water content, reinforcing its importance as a key covariate in gut microbiome studies [83,84].

Together, these results highlight that both environmental context and host-related factors shape gut microbiome variation in chacma baboons.

### Anthropization as a major driver of taxonomic and functional shifts

Anthropogenic disturbances, measured by HFI, strongly shaped the baboon gut microbiome by reducing alpha diversity. These results indicate that anthropogenic disturbance is a major driver of gut microbiome restructuring in chacma baboons, consistent with findings in humans [85–89] and wildlife [35,36,42], likely reflecting dietary shifts, particularly reduced fiber intake, coupled with limited exposure to environmental microbes caused by reduced home ranges in anthropized environments [90]. HFI also markedly restructured microbial composition, with effects robust to spatial variation. Additionally, no difference in within-troop variance was detected, suggesting directional shifts across individuals exposed to similar environmental conditions, rather than increased heterogeneity.

Taxonomically, HFI was associated with a decline in fiber-degradating taxa (Lachnospiraceae and Ruminococcaceae) and an increase with simple carbohydrate digestion (Lactobacillaceae) and dairy digestion (*Bifidobacterium*) taxa. These shifts mirror patterns described in industrialized human populations [85–89] and urban wildlife exploiting energy-rich human-derived food [41,43,54,91]. As Cape chacma baboons exploit comparable anthropogenic food resources [92], our findings suggest convergent microbial responses in primates exposed to dietary shifts. While often considered beneficial, the overabundance of *Bifidobacterium* and some Lactobacillaceae genera has been linked to obesity and metabolic dysregulation in industrialized human populations [93], although the implications for baboon health remain unclear.

However, some of our results contrasted with patterns reported in industrialized human populations. In humans, several genera such as *Prevotella, Treponema*, and *Succinivibrio*, appear depleted in urban context as compared to non-industrialized lifestyles [86,88,94]. In contrast, in chacma baboons, we observed an increase in *Treponema* and *Succinivibrio* with HFI, while *Prevotella* showed no change. Such differences likely reflect species-specific ecological contexts [36,42,43,54]. *Treponema* represents a diverse genus including species with distinct ecological functions. Baboons adopt a mixed foraging strategy, with continued access to natural foods, which may sustain or even favour certain taxa, such as *Treponema*, rather than leading to their loss as observed in industrialized human diets. An increase in *Succinivibrio* has also been reported in baboons with unlimited access to human waste [54], reflecting a metabolic response to energy-rich anthropogenic diets, although the underlying mechanisms require further investigation. The lack of *Prevotella* variation may be due to variation and functional differences within that genus, as two increasing ASVs could have partially offset the decrease of six others, suggesting a taxonomic restructuring rather than a total loss.

Functional shifts mirrored these taxonomic patterns, revealing a decline of complex organic substrate degradation and favouring central carbohydrate metabolism. Surprisingly, degradation pathways of glyphosate, toluene, and salicylate decreased with HFI, despite likely higher exposure in anthropized areas. This may reflect the loss of microbial diversity, resulting in functional erosion as taxa involved in complex degradation are depleted, as found in colobus monkeys, where these pathways facilitate breakdown of toxic plant components found in natural forest [95]. The positive association between glyphosate degradation and cultivated land indicates that agricultural environments, rather than urban areas, represent the primary sites of herbicide exposure.

### Diverging microbial responses to specific land-use variables

Beyond HFI, land-use variables revealed contrasting microbial responses. Built-up areas showed patterns broadly consistent with HFI, including reduced microbial diversity and a shift from fiber-degrader to simple carbohydrate taxa, but with weaker effects. These results highlight that microbial community responses reflect the cumulative effects of multiple anthropogenic pressures rather than a single environmental factor.

In contrast, cultivated areas showed distinct microbial signatures, likely reflecting specific environmental conditions, such as access to crop resources, livestock proximity, and/or agrochemical exposure. Specifically, the enrichment of *Butyrivibrio, Campylobacter*, and *Anaerovibrio* may reflect a livestock-associated signature [96], from environmental spillover or ruminant-like diet. The depletion of *Ruminococcus* and *Bifidobacterium* could be linked to glyphosate exposure, leading to potential gut dysbiosis [97,98]. This divergence suggests that different types of anthropization exert distinct ecological pressures on the gut microbiome. Despite these differences, convergent signals emerged: the depletion of *Parabacteroides* across HFI, built-up and cultivated, and an increase in *Bacteroides* in built-up and cultivated areas. These shifts indicate a shared loss of anti-inflammatory taxa and an increase of animal protein/fat rich diet [86,99].

Together, these results reinforce that different components of anthropization exert non-uniform effects on the gut microbiome, driving distinct and sometimes contrasting microbial responses depending on the type and intensity of anthropogenic disturbance.

### Isotopic dietary markers captured limited but specific microbial shifts

Dietary differences inferred from stable isotopes had comparatively limited effect on the baboon gut microbiome, showing no detectable impact on diversity and being associated with a relatively small subset of microbial taxa compared with broader anthropogenic variables. This suggests that anthropogenic environments may shape microbiome composition through multiple ecological pathways beyond diet alone. Dietary effects were weaker than those detected in other studies. Moy et al. [54] reported reduced microbial diversity in baboons with unrestricted access to human food waste, whereas baboons with limited or occasional access exhibited microbiomes similar to troops without access to trash. This contrast suggests that dietary changes in our study were either insufficient in magnitude, or stable isotopes failed to capture the specific nutritional signals - such as fiber or fat intake - that typically drive microbial richness and composition in anthropogenic areas [24,43,100]. In addition, landscape-level variables (e.g. built-up and cultivated land use) may already capture part of the dietary variation, thereby reducing the apparent independent effect of isotopic predictors.

Overall, these results suggest that gut microbial community responses are primarily driven by shifts in the relative abundances of existing taxa, with reductions in commensal and increases in potentially pathogenic taxa, more than the acquisition of novel or loss of microbial members.

### Perspectives

This study relied on 16S rRNA sequencing, future work using shotgun metagenomics would enable a more detailed characterization of microbial functional potential and its responses to anthropization. A substantial proportion of the community variation remained unexplained, likely reflecting unmeasured ecological and host-related dimensions including pollutants, behavioural heterogeneity in resource use, and life-history variation (age, sex, and reproductive status). Future longitudinal and seasonal designs will be essential to fully resolve the complex microbial responses to anthropogenic environments.

### Conclusion

Our study demonstrates that anthropization strongly shapes gut microbiome diversity, composition, and functional potential in chacma baboons, with partial convergence toward human patterns. Our multi-metric approach reveals that gut microbial responses vary across different anthropogenic pressures, emphasizing the need for precise definition and quantification of habitat disturbance. While microbial plasticity may facilitate short-term benefits, such as anthropogenic resource exploitation, the resulting loss of commensal taxa and enrichment of microbes linked to metabolic disorders may compromise host health and increase pathogen susceptibility. More broadly, this study underscores the importance of integrating ecological, behavioural, and microbiome approaches to understand wildlife responses to anthropogenic changes. Extending such an interdisciplinary approach across species and environments is essential for predicting the ecological and evolutionary consequences of host–microbe interactions for wildlife and public health in a changing world.

## Acknowledgments

We are grateful to the Nelson Mandela University, SANPark, CapeNature, the municipalities involved (George, Knysna, Bitou, Overstrand and Cape Town), and all relevant authorities for providing permits, access, and logistical support throughout this study. We sincerely thank the field assistants and students who contributed to the sample’s collection: Charlize Jones, Julia Roberts, Jade Norton, Alexis Watson, Tayla van Heerden, Zwelakhe Nkuna, Soanna Dany, Anne Guionneau, and Nicholas Taylor Van Rooyen. We also warmly thank everyone that help with fieldwork, including local inhabitants, baboon monitors, SANParks rangers, and private landowners who shared valuable information on baboon sightings or granted access to their properties. Data presented in this publication were partly produced through the GenSeq technical facilities of MEEB (CNRS and University of Montpellier) hosted by ISEM (CNRS, University of Montpellier and IRD). We thank the laboratories and platforms involved in sample processing, and analyses, including the Inqaba Biotechnical Industries, and the Stable Light Isotope Lab (University of Cape Town). We especially thank Julie Luyt and Patricia Groenewald for their technical support. Finally, we thank Philippe Veber for his support in statistical analysis.

## Author contributions

C.L., B.R., F.P., L.S. and V.R. designed the study. V.R. acquired the fundings for the project. C.L. and A.R. collected samples and C.L. conducted the laboratory work. C.L. performed the bioinformatic processing and data analysis, under the supervision of B.R., F.P., L.S., and V.R. C.L. wrote the original draft of the manuscript. All authors reviewed and contributed to the manuscript and approved the final version.

## Data availability

Sequencing data are available in the NCBI Sequence Read Archive (PRJNA1445581). Bioinformatic and statistical scripts are available on GitHub (available upon publication), and the ASV table, associated metadata, taxonomic assignments and phylogenetic tree are available via zenodo repository (10.5281/zenodo.20374880).

## References

1. Martínez-Fernández J, Ruiz-Benito P, Zavala MA. Recent land cover changes in Spain across biogeographical regions and protection levels: Implications for conservation policies. Land Use Policy 2015;44:62–75. 10.1016/j.landusepol.2014.11.021.

2. Grimm NB, Faeth SH, Golubiewski NE et al. Global Change and the Ecology of Cities. Science 2008;319(5864):756–60. 10.1126/science.1150195.

3. Lewis SL, Maslin MA. Defining the Anthropocene. Nature 2015;519(7542):171–80. 10.1038/nature14258.

4. Haddad NM, Brudvig LA, Clobert J et al. Habitat fragmentation and its lasting impact on Earth’s ecosystems. Sci Adv 2015;1(2):e1500052. 10.1126/sciadv.1500052.

5. Gao H, Chen J, Wang B et al. A study of air pollution of city clusters. Atmos Environ 2011;45(18):3069–77. 10.1016/j.atmosenv.2011.03.018.

6. Williams NSG, Mcdonnell MJ, Phelan GK et al. Range expansion due to urbanization: Increased food resources attract Grey-headed Flying-foxes (Pteropus poliocephalus) to Melbourne. Austral Ecol 2006;31(2):190–8. 10.1111/j.1442-9993.2006.01590.x.

7. Szulkin M, Garroway CJ, Corsini M et al. How to Quantify Urbanization When Testing for Urban Evolution? In: Szulkin M, Munshi-South J, Charmantier A (eds), Urban Evolutionary Biology, 1st edn. n.p.: Oxford University PressOxford, 2020, 13–35. 10.1093/oso/9780198836841.003.0002.

8. Nyhus PJ. Human–Wildlife Conflict and Coexistence. Annu Rev Environ Resour 2016;41(1):143–71. 10.1146/annurev-environ-110615-085634.

9. Ripple WJ, Chapron G, López-Bao JV et al. Saving the World’s Terrestrial Megafauna. BioScience 2016;66(10):807–12. 10.1093/biosci/biw092.

10. Hulme-Beaman A, Dobney K, Cucchi T et al. An Ecological and Evolutionary Framework for Commensalism in Anthropogenic Environments. Trends Ecol Evol 2016;31(8):633–45. 10.1016/j.tree.2016.05.001.

11. Gallo T, Fidino M, Lehrer EW et al. Urbanization alters predator-avoidance behaviours. J Anim Ecol 2019;88(5):793–803. 10.1111/1365-2656.12967.

12. Schulte-Hostedde AI, Mazal Z, Jardine CM et al. Enhanced access to anthropogenic food waste is related to hyperglycemia in raccoons (Procyon lotor). Conserv Physiol 2018;6(1):coy026. 10.1093/conphys/coy026.

13. Bateman PW, Fleming PA. Big city life: carnivores in urban environments. J Zool 2012;287(1):1–23. 10.1111/j.1469-7998.2011.00887.x.

14. Benmazouz I, Jokimäki J, Lengyel S et al. Corvids in Urban Environments: A Systematic Global Literature Review. 2021. http://hdl.handle.net/2437/328620 (26 Nov. 2025, date last accessed).

15. Lyons J, Mastromonaco G, Edwards DB et al. Fat and happy in the city: Eastern chipmunks in urban environments. Behav Ecol 2017;28(6):1464–71. 10.1093/beheco/arx109.

16. Scheun J, Greeff D, Nowack J. Urbanisation as an important driver of nocturnal primate sociality. Primates 2019;60(4):375–81. 10.1007/s10329-019-00725-0.

17. Diamant ES, Yeh PJ. Complex patterns of morphological diversity across multiple populations of an urban bird species. Evolution 2024;78(7):1325–37. 10.1093/evolut/qpae067.

18. Santos EG, Pompermaier VT, Wiederhecker HC et al. Urbanisation-induced changes in the morphology of birds from a tropical city. Emu-Austral Ornithol 2023;123(4):291–302. 10.1080/01584197.2023.2253836.

19. Chowdhury S, Brown J, Swedell L. Anthropogenic effects on the physiology and behaviour of chacma baboons in the Cape Peninsula of South Africa. Conserv Physiol 2020;8(1):coaa066. 10.1093/conphys/coaa066.

20. Murray MH, Sánchez CA, Becker DJ et al. City sicker? A meta-analysis of wildlife health and urbanization. Front Ecol Environ 2019;17(10):575–83.

21. Soto-Calderón ID, Acevedo-Garcés YA, Álvarez-Cardona J et al. Physiological and parasitological implications of living in a city: the case of the white-footed tamarin (Saguinus leucopus). Am J Primatol 2016;78(12):1272–81. 10.1002/ajp.22581.

22. Moll RJ, Cepek JD, Lorch PD et al. What does urbanization actually mean? A framework for urban metrics in wildlife research. J Appl Ecol 2019;56(5):1289–300. 10.1111/1365-2664.13358.

23. Cryan JF, O’Riordan KJ, Cowan CSM et al. The Microbiota-Gut-Brain Axis. Physiol Rev 2019;99(4):1877–2013. 10.1152/physrev.00018.2018.

24. Newman TM, Shively CA, Register TC et al. Diet, obesity, and the gut microbiome as determinants modulating metabolic outcomes in a non-human primate model. Microbiome 2021;9(1):100. 10.1186/s40168-021-01069-y.

25. Visconti A, Le Roy CI, Rosa F et al. Interplay between the human gut microbiome and host metabolism. Nat Commun 2019;10(1):4505. 10.1038/s41467-019-12476-z.

26. Amato KR, Leigh SR, Kent A et al. The role of gut microbes in satisfying the nutritional demands of adult and juvenile wild, black howler monkeys (Alouatta pigra). Am J Phys Anthropol 2014;155(4):652–64. 10.1002/ajpa.22621.

27. Leshem A, Liwinski T, Elinav E. Immune-Microbiota Interplay and Colonization Resistance in Infection. Mol Cell 2020;78(4):597–613. 10.1016/j.molcel.2020.03.001.

28. Vuong HE, Pronovost GN, Williams DW et al. The maternal microbiome modulates fetal neurodevelopment in mice. Nature 2020;586(7828):281–6. 10.1038/s41586-020-2745-3.

29. Archie EA, Tung J. Social behavior and the microbiome. Curr Opin Behav Sci The integrative study of animal behavior, 2015;6:28–34. 10.1016/j.cobeha.2015.07.008.

30. Clayton JB, Gomez A, Amato K et al. The gut microbiome of nonhuman primates: Lessons in ecology and evolution. Am J Primatol 2018;80(6):e22867. 10.1002/ajp.22867.

31. Dong TS, Gupta A. Influence of Early Life, Diet, and the Environment on the Microbiome. Clin Gastroenterol Hepatol 2019;17(2):231–42. 10.1016/j.cgh.2018.08.067.

32. Alberdi A, Aizpurua O, Bohmann K et al. Do Vertebrate Gut Metagenomes Confer Rapid Ecological Adaptation? Trends Ecol Evol 2016;31(9):689–99. 10.1016/j.tree.2016.06.008.

33. Moeller AH, Sanders JG. Roles of the gut microbiota in the adaptive evolution of mammalian species. Philos Trans R Soc B Biol Sci 2020;375(1808):20190597. 10.1098/rstb.2019.0597.

34. Zhu L, Wang J, Bahrndorff S. Editorial: The Wildlife Gut Microbiome and Its Implication for Conservation Biology. Front Microbiol 2021;12:697499. 10.3389/fmicb.2021.697499.

35. Teyssier A, Rouffaer LO, Saleh Hudin N et al. Inside the guts of the city: Urban-induced alterations of the gut microbiota in a wild passerine. Sci Total Environ 2018;612:1276–86. 10.1016/j.scitotenv.2017.09.035.

36. Wasimuddin, Malik H, Ratovonamana YR et al. Anthropogenic Disturbance Impacts Gut Microbiome Homeostasis in a Malagasy Primate. Front Microbiol 2022;13. https://www.frontiersin.org/articles/10.3389/fmicb.2022.911275 (10 Jan. 2024, date last accessed).

37. Alpízar P, Risely A, Tschapka M et al. Agricultural Fast Food: Bats Feeding in Banana Monocultures Are Heavier but Have Less Diverse Gut Microbiota. Front Ecol Evol 2021;9. https://www.frontiersin.org/articles/10.3389/fevo.2021.746783 (31 Oct. 2023, date last accessed).

38. Sugden S, Sanderson D, Ford K et al. An altered microbiome in urban coyotes mediates relationships between anthropogenic diet and poor health. Sci Rep 2020;10(1):art. 1. 10.1038/s41598-020-78891-1.

39. Martínez-Mota R, Righini N, Mallott EK et al. Environmental Stress and the Primate Microbiome: Glucocorticoids Contribute to Structure Gut Bacterial Communities of Black Howler Monkeys in Anthropogenically Disturbed Forest Fragments. Front Ecol Evol 2022;10. https://www.frontiersin.org/articles/10.3389/fevo.2022.863242 (23 Nov. 2023, date last accessed).

40. Adair MG, Tolley KA, Vuuren BJ van et al. Anthropogenic reverberations on the gut microbiome of dwarf chameleons (Bradypodion). PeerJ 2025;13:e18811. 10.7717/peerj.18811.

41. Lee W, Hayakawa T, Kiyono M et al. Gut microbiota composition of Japanese macaques associates with extent of human encroachment. Am J Primatol 2019;81(12):e23072. 10.1002/ajp.23072.

42. Barelli C, Albanese D, Stumpf RM et al. The Gut Microbiota Communities of Wild Arboreal and Ground-Feeding Tropical Primates Are Affected Differently by Habitat Disturbance. mSystems 2020;5(3):http://doi.org/10.1128/msystems.00061-20. 10.1128/msystems.00061-20.

43. Amato KR, Back JP, Sardaro MLS et al. Supplementation With Human Foods Affects the Gut Microbiota of Wild Howler Monkeys. Am J Primatol 2025;87(4):e70029. 10.1002/ajp.70029.

44. Nguyen HK, Jones PJ, Kendal D et al. Wildlife Microbiomes and the City: A Systematic Review of Urban Impacts on Wildlife Bacterial Communities, Microbiota and Host. published online 1 Feb. 2024. 10.1530/MAH-24-0003.

45. White J, Amato KR, Decaestecker E et al. Editorial: Impact of anthropogenic environmental changes on animal microbiomes. Front Ecol Evol 2023;11. https://www.frontiersin.org/articles/10.3389/fevo.2023.1204035 (31 Oct. 2023, date last accessed).

46. Johnson C, Piel A, Forman D et al. The ecological determinants of baboon troop movements at local and continental scales. Mov Ecol 2015;3. 10.1186/s40462-015-0040-y.

47. Schreier AL, Schlaht RM, Swedell L. Meat eating in wild hamadryas baboons: Opportunistic trade-offs between insects and vertebrates. Am J Primatol 2019;81(7):e23029. 10.1002/ajp.23029.

48. Hoffman T, O’Riain JM. The Spatial Ecology of Chacma Baboons (Papio ursinus) in a Human-modified Environment. Int J Primatol 2011;32:308–28. 10.1007/s10764-010-9467-6.

49. Kennedy Overton E, Bernard A, Renaud PC et al. Land use influences the diet of chacma baboons (Papio ursinus) in South Africa. Glob Ecol Conserv 26 July 2024:e03100. 10.1016/j.gecco.2024.e03100.

50. Mazué F, Guerbois C, Fritz H et al. Less bins, less baboons: reducing access to anthropogenic food effectively decreases the urban foraging behavior of a troop of chacma baboons (Papio hamadryas ursinus) in a peri-urban area. Primates J Primatol 2023;64(1):91–103. 10.1007/s10329-022-01032-x.

51. Birnie-Gauvin K, Peiman KS, Gallagher AJ et al. Sublethal consequences of urban life for wild vertebrates. Environ Rev 2016;24(4):416–25. 10.1139/er-2016-0029.

52. Björk JR, Dasari MR, Roche K et al. Synchrony and idiosyncrasy in the gut microbiome of wild baboons. Nat Ecol Evol 2022;6(7):955–64. 10.1038/s41559-022-01773-4.

53. Grieneisen LE, Livermore J, Alberts S et al. Group Living and Male Dispersal Predict the Core Gut Microbiome in Wild Baboons. Integr Comp Biol 2017;57(4):770–85. 10.1093/icb/icx046.

54. Moy M, Diakiw L, Amato KR. Human-influenced diets affect the gut microbiome of wild baboons. Sci Rep 2023;13(1):part. 1. 10.1038/s41598-023-38895-z.

55. Bambi M, Galla G, Donati C et al. Gut microbiota variations in wild yellow baboons (Papio cynocephalus) are associated with sex and habitat disturbance. Sci Rep 2024;14(1):869. 10.1038/s41598-023-50126-z.

56. Tsukayama P, Boolchandani M, Patel S et al. Characterization of Wild and Captive Baboon Gut Microbiota and Their Antibiotic Resistomes. mSystems 2018;3(3):10.1128/msystems.00016-18. http://doi.org/10.1128/msystems.00016-18.

57. Venter O, Sanderson EW, Magrach A et al. Sixteen years of change in the global terrestrial human footprint and implications for biodiversity conservation. Nat Commun 2016;7(1):12558. 10.1038/ncomms12558.

58. Amarenco G. Bristol Stool Chart□: étude prospective et monocentrique de «□l’introspection fécale□» chez des sujets volontaires. Prog En Urol 2014;24(11):708–13. 10.1016/j.purol.2014.06.008.

59. Gassert F, Venter O, Watson JEM et al. An operational approach to near real time global high resolution mapping of the terrestrial Human Footprint. Front Remote Sens 2023;4. 10.3389/frsen.2023.1130896.

60. Slater K, Barrett A, Brown LR. Home range utilization by chacma baboon (Papio ursinus) troops on Suikerbosrand Nature Reserve, South Africa. PLOS ONE 2018;13(3):e0194717. 10.1371/journal.pone.0194717.

61. Department of Forestry, Fisheries and the Environment. South African National Land Cover Geospatial Data. 2022. https://www.dffe.gov.za/egis (4 June 2024, date last accessed).

62. Sirko W, Kashubin S, Ritter M et al. Continental-Scale Building Detection from High Resolution Satellite Imagery, 2107.12283. Preprint, arXiv, 29 July 2021. 10.48550/arXiv.2107.12283.

63. O’Brien DM. Stable Isotope Ratios as Biomarkers of Diet for Health Research. Annu Rev Nutr 2015;35:565–94. 10.1146/annurev-nutr-071714-034511.

64. Codron D, Lee-Thorp JA, Sponheimer M et al. Inter-and intrahabitat dietary variability of chacma baboons (Papio ursinus) in South African savannas based on fecal δ13 C, δ15 N, and %N. Am J Phys Anthropol 2006;129(2):204–14. 10.1002/ajpa.20253.

65. Gohl DM, Vangay P, Garbe J et al. Systematic improvement of amplicon marker gene methods for increased accuracy in microbiome studies. Nat Biotechnol 2016;34(9):942–9. 10.1038/nbt.3601.

66. Cadamuro VC, Bouakaze C, Croze M et al. Determined about sex: Sex-testing in 45 primate species using a 2Y/1X sex-typing assay. Forensic Sci Int Genet 2015;14:96–107. 10.1016/j.fsigen.2014.09.010.

67. Bolyen E, Rideout JR, Dillon MR et al. Reproducible, interactive, scalable and extensible microbiome data science using QIIME 2. Nat Biotechnol 2019;37(8):852–7. 10.1038/s41587-019-0209-9.

68. Callahan BJ, McMurdie PJ, Rosen MJ et al. DADA2: High-resolution sample inference from Illumina amplicon data. Nat Methods 2016;13(7):581–3. 10.1038/nmeth.3869.

69. McDonald D, Jiang Y, Balaban M et al. Greengenes2 unifies microbial data in a single reference tree. Nat Biotechnol 2024;42(5):715–8. 10.1038/s41587-023-01845-1.

70. R Core Team. R: A Language and Environment for Statistical Computing. R Foundation for Statistical Computing, 2025. https://www.R-project.org/.

71. Wickham H. Ggplot2: Elegant Graphics for Data Analysis, with Sievert C. Use R! Second edition, Cham: Springer international publishing, 2016.

72. McMurdie PJ, Holmes S. phyloseq: An R Package for Reproducible Interactive Analysis and Graphics of Microbiome Census Data. PLoS ONE 2013;8(4):e61217. 10.1371/journal.pone.0061217.

73. Oksanen J, Simpson GL, Blanchet FG et al. Vegan: Community Ecology Package, version 2. 8–0. 2025. https://vegandevs.github.io/vegan/.

74. Bates D, Mächler M, Bolker B et al. Fitting Linear Mixed-Effects Models Using lme4. J Stat Softw 2015;67(1). 10.18637/jss.v067.i01.

75. Hartig F. DHARMa: Residual Diagnostics for Hierarchical (Multi-Level / Mixed) Regression Models. 2025. https://github.com/florianhartig/dharma.

76. Dray S, Bauman D, Blanchet G et al. {adespatial}: Multivariate Multiscale Spatial Analysis. 2025.

77. Nearing JT, Douglas GM, Hayes MG et al. Microbiome differential abundance methods produce different results across 38 datasets. Nat Commun 2022;13(1):342. 10.1038/s41467-022-28034-z.

78. Benjamini Y, Hochberg Y. Controlling the False Discovery Rate: A Practical and Powerful Approach to Multiple Testing. J R Stat Soc Ser B Stat Methodol 1995;57(1):289–300. 10.1111/j.2517-6161.1995.tb02031.x.

79. Douglas GM, Maffei VJ, Zaneveld JR et al. PICRUSt2 for prediction of metagenome functions. Nat Biotechnol 2020;38(6):685–8. 10.1038/s41587-020-0548-6.

80. Lladó S, López-Mondéjar R, Baldrian P. Forest Soil Bacteria: Diversity, Involvement in Ecosystem Processes, and Response to Global Change. Microbiol Mol Biol Rev MMBR 2017;81(2):e00063–16. 10.1128/MMBR.00063-16.

81. Krishnappa C, Mahanta DK, Pandey S et al. Trends and patterns in microbial research within forest ecosystems: bibliometric insights from 2010 to 2025. Discov For 2026;2(1):2. 10.1007/s44415-025-00067-4.

82. Kim YS, Unno T, Kim BY et al. Sex Differences in Gut Microbiota. World J Mens Health 2020;38(1):48–60. 10.5534/wjmh.190009.

83. Procházková N, Falony G, Dragsted LO et al. Advancing human gut microbiota research by considering gut transit time. Gut 2023;72(1):180–91. 10.1136/gutjnl-2022-328166.

84. Vandeputte D, Falony G, Vieira-Silva S et al. Stool consistency is strongly associated with gut microbiota richness and composition, enterotypes and bacterial growth rates, Gut Microbiota. Gut 2016;65(1):57–62. 10.1136/gutjnl-2015-309618.

85. Rosas-Plaza S, Hernández-Terán A, Navarro-Díaz M et al. Human Gut Microbiome Across Different Lifestyles: From Hunter-Gatherers to Urban Populations. Front Microbiol 2022;13:843170. 10.3389/fmicb.2022.843170.

86. Sonnenburg ED, Sonnenburg JL. The ancestral and industrialized gut microbiota and implications for human health. Nat Rev Microbiol 2019;17(6):383–90. 10.1038/s41579-019-0191-8.

87. Abjani F, Madhavan P, Chong PP et al. Urbanisation and its associated factors affecting human gut microbiota: where are we heading to? Ann Hum Biol 2023;50(1):137–47. 10.1080/03014460.2023.2170464.

88. Vinogradova E, Mukhanbetzhanov N, Nurgaziyev M et al. Impact of urbanization on gut microbiome mosaics across geographic and dietary contexts. mSystems 2024;9(10):e00585–24. 10.1128/msystems.00585-24.

89. Zhang F, Zhou G, Schewe M et al. Dietary urbanization destabilizes host-gut microbiome homeostasis and informs precision nutrition for human health. Cell Metab 2025;37(11):2128–48. 10.1016/j.cmet.2025.09.013.

90. Hoffman TS, O’Riain MJ. Troop Size and Human-Modified Habitat Affect the Ranging Patterns of a Chacma Baboon Population in the C ape P eninsula, S outh A frica. Am J Primatol 2012;74(9):853–63. 10.1002/ajp.22040.

91. Li Y, Yan Y, Wu H et al. The role of gut microbiota in a generalist, golden snub-nosed monkey, adaptation to geographical diet change. Anim Microbiome 2024;6(1):part. 1. 10.1186/s42523-024-00349-w.

92. Doorn AC van, O’Riain MJ. Nonlethal management of baboons on the urban edge of a large metropole. Am J Primatol 2020;82(8):e23164. 10.1002/ajp.23164.

93. Chakraborti CK. New-found link between microbiota and obesity. World J Gastrointest Pathophysiol 2015;6(4):110–9. 10.4291/wjgp.v6.i4.110.

94. Tamburini FB, Maghini D, Oduaran OH et al. Short-and long-read metagenomics of urban and rural South African gut microbiomes reveal a transitional composition and undescribed taxa. Nat Commun 2022;13(1):926. 10.1038/s41467-021-27917-x.

95. Barelli C, Albanese D, Donati C et al. Habitat fragmentation is associated to gut microbiota diversity of an endangered primate: implications for conservation. Sci Rep 2015;5(1):part. 1. 10.1038/srep14862.

96. Forcina G, Pérez-Pardal L, Carvalheira J et al. Gut Microbiome Studies in Livestock: Achievements, Challenges, and Perspectives. Animals 2022;12(23). 10.3390/ani12233375.

97. Lehman PC, Cady N, Ghimire S et al. Low-dose glyphosate exposure alters gut microbiota composition and modulates gut homeostasis. Environ Toxicol Pharmacol 2023;100:104149. 10.1016/j.etap.2023.104149.

98. Walsh L, Hill C, Ross RP. Impact of glyphosate (RoundupTM) on the composition and functionality of the gut microbiome. Gut Microbes 2023;15(2):2263935. 10.1080/19490976.2023.2263935.

99. Singh RK, Chang HW, Yan D et al. Influence of diet on the gut microbiome and implications for human health. J Transl Med 2017;15(1):73. 10.1186/s12967-017-1175-y.

100. Lee W, Hayakawa T, Kiyono M et al. Diet-related factors strongly shaped the gut microbiota of Japanese macaques. Am J Primatol 2023;85(12):e23555. 10.1002/ajp.23555.

