## Supplementary material for "Human footprints in the gut: how anthropogenic environments reshape the microbiome of chacma baboons"

### Table of content


**Supplementary Materials and Methods**

**Supplementary results**

### Supplementary Tables

**Supplementary Table 1.** Characteristics of the studied chacma baboon troops, including sampling effort and quantitative measures of anthropogenic pressure and habitat composition across sites

**Supplementary Table 2.** Pairwise Bray–Curtis dissimilarity and sequencing depth of technical replicates sequenced across two independent MiSeq runs, used to assess sequencing reproducibility and batch effects.

**Supplementary Table 3.** Results of PERMANOVA models based on weighted and unweighted UniFrac distances testing shifts in gut microbial community composition

**Supplementary Table 4.** Results of dbRDA models based on weighted and unweighted UniFrac distances, testing shifts in gut microbial community composition while accounting for spatial structure

**Supplementary Table 5.** Linear mixed model results predicting variation in gut microbial community dispersion across baboon troops.

**Supplementary Table 6.** ANCOM-BC results of taxonomic and functional differential abundance in chacma baboon along different anthropogenic variables.

### Supplementary Figures

**Supplementary Figure 1.** Comparison of ASV composition between soil and fecal samples and assessment of potential environmental contamination.

**Supplementary Figure 2.** Rarefaction curves and beta-diversity structure used to define sequencing depth threshold and identify duplicates.

**Supplementary Figure 3.** Bioinformatic filtering workflow and sample/sequence retention across processing steps.

**Supplementary Figure 4.** Relationships between stable isotope values and gut microbial alpha diversity, illustrating the decomposition of $\delta^{15}N$ effects.

**Supplementary Figure 5.** Distribution of stable isotope values ($\delta^{13}C$ and $\delta^{15}N$) across samples along the anthropogenic gradient.

**Supplementary Figure 6.** Pairwise Spearmann correlations among all variables considered prior the analysis.

**Supplementary Figure 7.** Taxonomic composition of the gut microbiome across baboon troops along the anthropogenic gradient.

**Supplementary Figure 8.** Difference CAPE/GAR alpha, beta diversity and differential abundance.

**Supplementary Figure 9.** Sexe difference in alpha, beta diversity and differential abundance in baboons.

**Supplementary Figure 10.** Overlap of differentially abundant genera across anthropogenic variables.

### Supplementary Materials and Methods

#### Sample collection

A total of 512 fecal samples were collected non-invasively from 33 troops of chacma baboons between March and May 2024. Sampling was conducted within a narrow temporal window to minimize seasonal variation in diet and microbiome composition [1,2]. Only fresh fecal samples were collected, and samples likely originating from infants were excluded to avoid age-related variation in gut microbiota [3,4]. Samples were collected from the internal portion of the feces to minimize external contamination and homogenised. Each sample was split into two subsamples: one preserving 5g of feces in RNAlater (Thermo Fisher Scientific) for microbiome analysis and one of 5g without preservative for stable isotope analysis. For each sample, we recorded the GPS location, date, time, and Bristol Stool Scale (see Data availability). The Bristol Stool Scale classifies feces based on shape and consistency, ranging from hard lumps (1) to entirely liquid stools (7) [5]. To control for potential environmental contamination, 52 soil samples or swabs were collected in proximity to fecal samples across substrate types. All samples were kept at 4°C during fieldwork, and transferred to -20°C within 12h of collection at Nelson Mandela University Molecular laboratory until further processing.

#### Anthropization characterization

HFI values were downloaded rescaled (x 1,000) in uint16 data type to reduce file sizes (e.g. 50 000 corresponds to an HFI of 50).

Baboon diet was characterized using stable isotope analysis of nitrogen (δ15N) and carbon (δ13C) from fecal samples. Samples were prepared following standard protocols [6]. Briefly, fecal samples were thawed (4°C, 24h), oven-dried (60°C, 48h), homogenised, then ground and sieved through a 600µm mesh and ~0.9-1.2 mg of material was encapsulated (Elemtex Cat No: SE1003.250, UK) for analysis. All isotopic analyses were conducted at the Stable Light Isotope Laboratory, Archaeology Department, University of Cape Town. As part of quality control procedures, 5% of fecal samples (every 20th sample) were prepared in duplicates. Blank samples and blank capsules were also included in each run to assess contamination.

Isotopic measurements were performed using an elemental analyser (Flash 2000, Thermo Scientific), coupled to an isotope ratio mass spectrometer (Delta V Plus), operated via Isodat© 3.0 software. Analytic precision of the isotopic standards measured within the same run as samples was < 0.2‰ for both $\delta^{13}C$ and $\delta^{15}N$. Four internal standards calibrated against international reference material were used for the analysis of the fecal material (Acacia: $\delta^{13}C$ = -28.14‰, $\delta^{15}N$ = -0.7‰; Valine: $\delta^{13}C$ = -27.02‰, $\delta^{15}N$ = 12.33‰; MG New: $\delta^{13}C$ = -27.02‰, $\delta^{15}N$ = 12.33‰; and ANU Sucrose: $\delta^{13}C$ = -10.4‰). $\delta^{13}C$ values are expressed relative to Vienna Pee Dee Belemnite (VPDB); $\delta^{15}N$ values are expressed relative to atmospheric nitrogen (AIR).

#### DNA extraction and sequencing of the bacterial V4 16S rRNA gene

Gut microbial communities were characterized from 512 fecal samples and 52 soil samples preserved in RNAlater using the Qiagen DNeasy PowerSoil Pro kit (QIAGEN, Germany), following the manufacturer protocol with a bead-beating step for cell lysis. The V4 hypervariable region of the 16S rRNA gene was amplified using the primers 515F (5'-GTGYCAGCMGCCGCGGTAA-3') and 806R (5'-GGACTACNVGGGTWTCTAAT-3') by PCR, following the dual indexing protocol described in [7]. Primers included Illumina adapter overhangs for downstream indexing. Extraction blanks (39), PCR negative controls (39) and sequencing blanks (9) were included to monitor potential contamination and were processed in parallel with fecal and soil samples. All PCR reactions were performed in duplicate to minimize potential amplification biases. After verification of the PCR amplifications by electrophoresis on a 1.5% agarose gel, each sample's amplicon duplicates were pooled and sent to GenSeq laboratory (Montpellier, France) for further processing. The PCR amplicons were purified using SPRI bead-based cleanup (AMPure XP, Beckman Coulter), and a second indexing PCR was performed to attach Illumina dual indices and sequencing adapters. Equimolar amplicons were pooled and sequenced with two independent Illumina MiSeq runs (V3 chemistry, 600 cycles, 2 x 300 bp paired-end). To assess potential batch effects, ten samples were sequenced in duplicate across both runs.

### Bioinformatic processing

#### Read processing and taxonomy assignment

After sequencing, eight samples failing sequencing were removed, resulting in 504 fecal samples retained for analyses (plus 10 sequencing replicates). The mean sequencing depth of fecal samples before filtering was 57,736 reads per sample (range: 17,523 - 99,986). Sequences were processed using QIIME2 (v.2024.10) [8]. Primer sequences were removed using cut-adapt (cutadapt 2024.10.0) and reads were quality-filtered, denoised, merged, and chimera-checked using DADA2, generating an Amplicon Sequence Variants (ASVs) table [9]. Forward and reverse reads were trimmed to 240 and 160 bases, respectively, based on the phred quality score distribution (Q>30), to remove low-quality regions. Each run was processed individually and merged prior to downstream analyses. After denoising, 23,801,111 reads were retained (mean 46,396 reads per sample; range: 14,334 – 73,034). Taxonomic assignment was performed Naive Bayes classifier trained on the GreenGenes2 database (2024.09) [10,11]. Mitochondrial and chloroplast sequences were removed, resulting in 6,417 ASVs. An unrooted phylogenetic tree was constructed using MAFFT for sequence alignment and FastTree for tree inference, implemented within the QIIME2 align-to-tree-mafft-fasttree pipeline, and midpoint-rooted. Resulting feature tables, taxonomy, and phylogenetic tree were imported into R version 4.5.2 using qiime2R v.0.99.6 for downstream analysis [12,13].

#### Control and sample filtration

Of the 87 negative controls (DNA extraction, PCR and sequencing blanks), 32 samples contained very low read counts after the denoising/filtering steps described above (29 ASVs in total; 1-3 reads per ASV), and the rest had no sequence at all. Among the 29 ASVs detected in controls, 19 also appeared in fecal samples, but at much higher abundance in feces suggesting minor cross-contaminations coming from fecal samples rather than laboratory contaminants. Given the negligible read counts, all negative controls were excluded from further analyses.

We then assessed sequencing batch effects between the two runs by comparing their gut microbial community similarity, using the beta-diversity (Bray–Curtis) calculated with the phyloseq package in R [14]. Sequencing replicates showed high similarity (Bray–Curtis < 0.25; Supplementary Table 2), and only the replicate with the highest read count was retained for each pair.

Potential soil contamination was evaluated by comparing ASVs overlap between fecal and soil samples at the troop level. Nine soil samples had no sequences left after the denoising steps, and we identified 7,998 ASVs across the remaining 43 soil samples (mean ± SD ASVs per sample: 278 ± 296, range: 1-29,447). Only 0–15 ASVs were shared between soil and feces per troop (146 ASVs in total), representing < 1% of fecal ASVs (Supplementary Fig. 1A-B). To be conservative, we removed any of these shared ASVs that were more abundant in soil than in feces based on either (i) cumulative or (ii) mean read counts, corresponding to a total of 57 ASVs (Supplementary Fig. 1C). Note that among these 57 ASVs, 50 would have been excluded by subsequent filtering steps removing low-abundance ASVs (< 50 reads). This approach ensured that only ASVs likely originating from environmental contamination were excluded while retaining biologically relevant taxa.

To verify sample identity, a subset of samples (n = 21) for which species assignment was uncertain in the field, was screened using mitochondrial 16S rRNA sequencing with universal mammalian primers (16Smam1 and 16Smam2) [15], followed by Sanger sequencing (Inqaba Biotechnical Industries, South Africa). This allowed the identification of five non-baboon samples (four *Sus scrofa* samples and one *Potamochoerus porcus* sample), which were excluded from the dataset.

In addition, pairwise beta-diversity (Bray-Curtis) was used to assess microbial similarity between samples. To compare the microbial composition across fecal samples, data were rarefied to 25,000 reads per sample, based on alpha rarefaction curves generated with the vegan package v2.7.1 in R (Supplementary Fig. 2A). The distribution of pairwise dissimilarities showed a dominant peak corresponding to inter-individual variation (Supplementary Fig. 2B), as well as a secondary peak of very low dissimilarity values indicative of duplicate samples from the same individual (Supplementary Fig. 2C). Based on this pattern, a threshold of 0.25 was applied to identify duplicate samples, and the sample with the highest read count was retained for each pair. This threshold was supported by the clear separation between low- and high-dissimilarity values (Supplementary Fig. 2D). This resulted in the removal of 51 duplicate samples, mostly collected on the same day, except two pairs collected one day apart.

After filtering, 448 fecal samples were retained, with a mean sequencing depth of 46,310 reads per sample (range: 13,524 - 72,359) and a total of 3,552 ASVs.

**Sex characterization**

Individual sex was determined using a multiplex PCR targeting the UTX, UTY and SRY regions, following the protocol described in [16]. Sex was assigned based on banding patterns after gel electrophoresis: females showed a single UTX band, whereas males showed two or three bands corresponding to X- and Y-linked markers.

**Statistical analyses**

All analyses were performed using R v4.5.2 [12] and figures were generated using ggplot2 [17] and edited with Inkscape. Anthropization was modelled using a two-step framework, including (i) the Human Footprint Index (HFI) and (ii) finer-scale variables combining land-use proportions and stable isotopes ($\delta^{15}N$ and $\delta^{13}C$).

**Environmental and host predictors of the gut microbiome alpha-diversity.** Alpha diversity was assessed using three metrics: observed richness (total number of unique ASVs per sample), Shannon Index (accounting for both richness and evenness in abundance of ASVs), and Faith's phylogenetic diversity (Faith's PD, integrating phylogenetic relationships). These metrics were calculated using the phyloseq v1.52.0 and vegan v2.7.1 packages [14,18]. To analyze the influence of anthropization on alpha-diversity, linear and generalized linear mixed models (LMMs/ GLMMs) [19] were fitted for each alpha diversity metrics using lme4 version 1.1.37 [19]. Observed species richness was modelled using a negative binomial error distribution, while Shannon diversity and Faith's PD were modelled using a Gaussian distribution. Following our two-step analytical framework, anthropization was modelled using either (1) the Human Footprint Index or (2) land-use proportions and stable isotope values ($\delta^{15}N$ and $\delta^{13}C$), as the fixed effects. $\delta^{15}N$ values varied both between troops and within troops, with opposite patterns observed at these two levels for alpha diversity (Supplementary Fig. 4A-B). To disentangle intra- and inter-troop variations, we decomposed $\delta^{15}N$ into two components: $\delta^{15}N$_mean, representing the mean value for each troop, and $\delta^{15}N$_diff, representing the deviation of each sample from its troop mean. No such divergence was observed for $\delta^{13}C$ (Supplementary Fig. 4C-D) and was retained as a single variable without decomposition. In addition, we included the sex and the Bristol Stool Scale as control variables, as there is evidence suggesting that these factors are associated with variation in gut microbiome diversity [20–22]. We also included the geographical area (GAR/CAPE) as a control variable to account for potential environmental differences, as well as the studied troop as a random effect to account for non-independence among samples. Variance inflation factors (VIF) were calculated using the car package v3.1.3 to assess collinearity among explanatory variables. Following this analysis, among the land-use variables, only built-up and cultivated proportions were retained in the models, as the others were either highly collinear (VIF > 2) or less informative, while these two variables best captured biologically relevant gradients of anthropogenic disturbance. Pairwise correlations among predictors are shown in Supplementary Fig. 6. Therefore, for approach (2), final models included built-up and cultivated proportions as land-use variables, stable isotope values ($\delta^{15}N\_mean$, $\delta^{15}N\_diff$ and $\delta^{13}C$), sex, Bristol Stool Scale and area, as fixed effects. To facilitate model convergence, all continuous variables were centered and scaled. We tested the interactions for models (1) between human footprint and area or sex, and for models (2) between sex and built-up or cultivated land-use variables. Assumptions of normality and homoscedasticity of residuals were checked with DHARMa package v0.4.7 [23], which generates simulated residuals, revealing no significant deviations. All possible model subsets were generated using the *dredge* function of the MuMIn package v1.48.11 [24], and model selection was based on the Akaike Information Criterion corrected for small sample size (AICc). Then, we calculated model-averaged coefficients across models with $\Delta$AICc < 2. Furthermore, we also tested for spatial autocorrelation between samples in the simulated scaled residuals of the fitted models with DHARMa package.

**Environmental and host predictors of the gut microbiome composition. .** Inter-individual variation in gut microbial community composition was assessed using weighted and unweighted UniFrac distances, which account for phylogenetic relatedness among taxa, and weighted UniFrac additionally accounting for relative abundances, using the phyloseq package [14]. First, we tested whether beta-diversity varied along the anthropogenic gradient using the permutational multivariate analysis of variance (PERMANOVA), implemented in the *adonis2* function of the vegan package [18] under two frameworks. The first model included the Human Footprint Index, sex and area as fixed variables, while the second model included the land-use variables (built-up and cultivated), the stable isotope variables ($\delta^{15}N$ and $\delta^{13}C$), sex and area. We implemented a restricted permutation procedure to account for non-independence among samples within troops. For each predictor, values were randomly reassigned among troops, such that all samples within a troop were assigned the same permuted value, and the PERMANOVA was rerun on each permuted dataset. This procedure was repeated 9,999 times to generate null distribution of F-statistics, against which observed F-value from the original (unpermuted) model were compared to assess significance. Second, we evaluated potential spatial autocorrelation of the beta diversity using a Mantel test between weighted and unweighted UniFrac dissimilarity matrices and geographical distance matrices (derived from troops' coordinates), with 'area' as the strata argument. Moreover, we investigated the correlations between weighted and unweighted UniFrac microbial dissimilarity matrices and the environmental variables, controlling for the effect of the geographical coordinates of the study troops using distance-based redundancy analysis (dbRDA) [25]. To avoid non-independence among samples, data were averaged at the troop level. Spatial structure was accounted for by including distance-based Moran's eigenvector maps (dbMEMs) as conditional variable. dbRDA analyses were performed using the *capscale* function in the vegan package [18], with spatial variables generated using the *dbmem* function in the adespatial package v.0.3-28 [26]. Statistical significance was assessed using permutation-based ANOVA with 9,999 permutations.

In addition, beta-diversity dispersion along the human footprint gradient was assessed using the PERMDISP2 test implemented in the *betadisper* function of the vegan package [18]. Distances to group centroids were calculated at the troop level to account for troop structure and to limit biases related to uneven sampling. To avoid small-sample size effects, only troops with more than 10 samples were included in the analysis. Distances to group centroids were then modelled using LMMs from the lme4 package [19] assuming a Gaussian distribution. The fixed effects included in the model were the Human Footprint Index, sex and area, while troop identity was included as a random effect.

**Microbial functional prediction.** Microbial functional profiles were predicted from 16S rRNA data (unrarefied dataset) using PICRUSt2 with default parameters [27]. ASVs with nearest-sequenced taxon index (NSTI) > 2, were excluded. Predicted functional profiles included Enzyme Commissions (EC) and MetaCyc pathways. Pathways assigned to plant-related (e.g. C4 photosynthetic carbon assimilation cycle and allantoin degradation in plants) were excluded. Differential abundance of functional pathways across the anthropization gradient was assessed using ANCOM-BC2 [28] under both frameworks, as for taxonomic analysis.

### References


1. Hicks AL, Lee KJ, Couto-Rodriguez M *et al.* Gut microbiomes of wild great apes fluctuate seasonally in response to diet. *Nat Commun* 2018;**9**(1):1786. https://doi.org/10.1038/s41467-018-04204-w.
2. Murillo T, Schneider D, Fichtel C *et al.* Dietary shifts and social interactions drive temporal fluctuations of the gut microbiome from wild redfronted lemurs. *ISME COMMUN* 2022;**2**(1):art. 1. https://doi.org/10.1038/s43705-021-00086-0.
3. Baniel A, Petrullo L, Mercer A *et al.* Maternal effects on early-life gut microbiota maturation in a wild nonhuman primate. *Current Biology* 2022;**32**(20):4508-4520.e6. https://doi.org/10.1016/j.cub.2022.08.037.
4. Reese AT, Phillips SR, Owens LA *et al.* Age patterning in wild chimpanzee gut microbiota diversity reveals differences from humans in early life. *Curr Biol* 2021;**31**(3):613-620.e3. https://doi.org/10.1016/j.cub.2020.10.075.
5. Amarenco G. Bristol Stool Chart : étude prospective et monocentrique de « l'introspection fécale » chez des sujets volontaires. *Progrès en Urologie* 2014;**24**(11):708–13. https://doi.org/10.1016/j.purol.2014.06.008.
6. Codron D, Lee-Thorp JA, Sponheimer M *et al.* Inter- and intrahabitat dietary variability of chacma baboons ( *Papio ursinus* ) in South African savannas based on fecal $\delta^{13}$ C, $\delta^{15}$ N, and %N. *American J Phys Anthropol* 2006;**129**(2):204–14. https://doi.org/10.1002/ajpa.20253.
7. Gohl DM, Vangay P, Garbe J *et al.* Systematic improvement of amplicon marker gene methods for increased accuracy in microbiome studies. *Nat Biotechnol* 2016;**34**(9):942–9. https://doi.org/10.1038/nbt.3601.
8. Bolyen E, Rideout JR, Dillon MR *et al.* Reproducible, interactive, scalable and extensible microbiome data science using QIIME 2. *Nat Biotechnol* 2019;**37**(8):852–7. https://doi.org/10.1038/s41587-019-0209-9.
9. Callahan BJ, McMurdie PJ, Rosen MJ *et al.* DADA2: High-resolution sample inference from Illumina amplicon data. *Nat Methods* 2016;**13**(7):581–3. https://doi.org/10.1038/nmeth.3869.
10. Bokulich NA, Kaehler BD, Rideout JR *et al.* Optimizing taxonomic classification of marker-gene amplicon sequences with QIIME 2's q2-feature-classifier plugin. *Microbiome* 2018;**6**(1):90. https://doi.org/10.1186/s40168-018-0470-z.
11. McDonald D, Jiang Y, Balaban M *et al.* Greengenes2 unifies microbial data in a single reference tree. *Nat Biotechnol* 2024;**42**(5):715–8. https://doi.org/10.1038/s41587-023-01845-1.
12. R Core Team. *R: A Language and Environment for Statistical Computing*. R Foundation for Statistical Computing, 2025. https://www.R-project.org/.
13. Bisanz JE. *qiime2R: Importing QIIME2 Artifacts and Associated Data into R Sessions*. 2018. https://github.com/jbisanz/qiime2R.
14. McMurdie PJ, Holmes S. phyloseq: An R Package for Reproducible Interactive Analysis and Graphics of Microbiome Census Data. *PLoS ONE* 2013;**8**(4):e61217. https://doi.org/10.1371/journal.pone.0061217.
15. Taylor PG. Reproducibility of ancient DNA sequences from extinct Pleistocene fauna. *Molecular Biology and Evolution* 1996;**13**(1):283–5. https://doi.org/10.1093/oxfordjournals.molbev.a025566.
16. Cadamuro VC, Bouakaze C, Croze M *et al.* Determined about sex: Sex-testing in 45 primate species using a 2Y/1X sex-typing assay. *Forensic Science International: Genetics* 2015;**14**:96–107. https://doi.org/10.1016/j.fsigen.2014.09.010.

17. Wickham H. *Ggplot2: Elegant Graphics for Data Analysis*, with Sievert C. Use R! Second edition, Cham: Springer international publishing, 2016.
18. Oksanen J, Simpson GL, Blanchet FG *et al. Vegan: Community Ecology Package*, version 2.8-0. 2025. https://vegandevs.github.io/vegan/.
19. Bates D, Mächler M, Bolker B *et al.* Fitting Linear Mixed-Effects Models Using **lme4**. *J Stat Soft* 2015;**67**(1). https://doi.org/10.18637/jss.v067.i01.
20. Bambi M, Galla G, Donati C *et al.* Gut microbiota variations in wild yellow baboons (Papio cynocephalus) are associated with sex and habitat disturbance. *Sci Rep* 2024;**14**(1):869. https://doi.org/10.1038/s41598-023-50126-z.
21. Kim YS, Unno T, Kim BY *et al.* Sex Differences in Gut Microbiota. *World J Mens Health* 2020;**38**(1):48–60. https://doi.org/10.5534/wjmh.190009.
22. Vandeputte D, Falony G, Vieira-Silva S *et al.* Stool consistency is strongly associated with gut microbiota richness and composition, enterotypes and bacterial growth rates, Gut Microbiota. *Gut* 2016;**65**(1):57–62. https://doi.org/10.1136/gutjnl-2015-309618.
23. Hartig F. *DHARMa: Residual Diagnostics for Hierarchical (Multi-Level / Mixed) Regression Models*. 2025. https://github.com/florianhartig/dharma.
24. Bartoń K. *MuMIn: Multi-Model Inference*. 2025. https://CRAN.R-project.org/package=MuMIn.
25. Legendre P, Fortin MJ, Borcard D. Should the Mantel test be used in spatial analysis? *Methods in Ecology and Evolution* 2015;**6**(11):1239–47. https://doi.org/10.1111/2041-210X.12425.
26. Dray S, Bauman D, Blanchet G *et al. {adespatial}: Multivariate Multiscale Spatial Analysis*. 2025.
27. Douglas GM, Maffei VJ, Zaneveld JR *et al.* PICRUSt2 for prediction of metagenome functions. *Nat Biotechnol* 2020;**38**(6):685–8. https://doi.org/10.1038/s41587-020-0548-6.
28. Nearing JT, Douglas GM, Hayes MG *et al.* Microbiome differential abundance methods produce different results across 38 datasets. *Nat Commun* 2022;**13**(1):342. https://doi.org/10.1038/s41467-022-28034-z.

### Supplementary results

#### Habitat characteristics, anthropogenic pressure, and diet across baboon troops

We studied 512 fecal samples from 33 chacma baboon troops distributed along an anthropogenic gradient in the Western Cape, South Africa (Fig. 1A-C). Sex was determined for most samples (317 females, 185 males, 10 unknown), with females representing on average 62% (range: 0-100%) of samples per troop.

Troops were distributed across two areas that differed in their natural habitat composition: the CAPE area was primarily characterized by shrubland (covering ~50%), whereas the GAR area was dominated by forested habitats (~70%) (Fig. 1A; Supplementary Table 1). Anthropogenic pressure spanned comparable ranges between areas, with HFI values ranging from 9,880 to 29,276 in CAPE and 5,700 to 29,281 in GAR (Supplementary Table 1). Land-use composition varied within both areas, reflecting environmental heterogeneity along the anthropogenic gradient. The mean proportion of built-up areas was 20% (range: 0-47%) in the CAPE area and 12% (range: 0-63%) in the GAR area. Conversely, the mean proportion of cultivated land was 4% (range: 0-19%) and 14% (0-54%) in the CAPE and GAR area, respectively (Fig. 1B-C). Overall, these values illustrate a gradient of landscape transformation in both areas.

Diet was characterized using stable isotope values (δ15N and δ13C). Across the two areas, the mean $\delta^{15}N$ value was 3.96‰ (range: –0.24 to 6.74‰), and the mean $\delta^{13}C$ value was –25.94‰ (range: –30.82 to –18.71‰) (Supplementary Fig. 5A). $\delta^{15}N$ values differed significantly between areas, with lower values observed in GAR compared to CAPE (LM, $\beta$ = -0.42 ± .011, t = -3.68, p<0.001), and increased with HFI (LM, $\beta$ = 0.21 ± 0.06, t = 3.63, p<0.001) (Supplementary Fig. 5B). No significant difference was detected in $\delta^{13}C$ with HFI (LM, p=0.87). No significant differences in stable isotope values were observed between sexes (LM, $\delta^{15}N$: p=0.59; $\delta^{13}C$: p=0.27) and Bristol Stool Scale was not associated with HFI (LM, p=0.66).

**Supplementary Table 1.** Characteristics of the studied chacma baboon troops, including sampling effort and quantitative measures of anthropogenic pressure and habitat composition across sites. For each troop, we report its geographic area (CAPE or GAR), estimated troop size (number of individuals), number of fecal samples collected, and the presence or absence of active baboon monitoring. Anthropogenic pressure is described using the Human Footprint Index (HFI), expressed as scaled values (×1,000; i.e. 50,000 = HFI 50), as well as the percentage of built-up and cultivated land within a 2 km² buffer around sampling locations. Dominant land cover corresponds to the main habitat type within each buffer, based on the South African National Land Cover dataset (2022).

| Troop number | Troop | Area | Troop size | Samples | Human Footprint | Built-up (%) | Cultivated (%) | Dominated land cover |
|---|---|---|---|---|---|---|---|---|
| **1** | Table Mountain National park - Zwaanswyk | CAPE | 32 | 17 | 19 403 | 15.3 | 16.3 | Forest |
| **2** | Table Mountain National park - Tokai | CAPE | 115 | 31 | 20 414 | 22.1 | 11.6 | Forest |
| **3** | Da Gama | CAPE | 20 | 15 | 24 283 | 23.0 | 0.0 | Shrubland |
| **4** | Simon's Town - Waterfall | CAPE | 42 | 22 | 29 276 | 46.7 | 0.0 | Shrubland |
| **5** | Simon's Town - Seaforth | CAPE | 16 | 3 | 24 071 | 37.7 | 0.0 | Shrubland |
| **6** | Cape of good Hope-Olifantsbos | CAPE | 46 | 14 | 11 709 | 0.0 | 0.0 | Shrubland |
| **7** | Cape pf good Hope – Buffel Bay | CAPE | 33 | 22 | 9 880 | 0.7 | 0.0 | Shrubland |
| **8** | Roo-Els | CAPE | 24 | 12 | 16 824 | 25.6 | 0.0 | Shrubland |
| **9** | Pringle Bay | CAPE | 21 | 10 | 20 584 | 41.5 | 0.0 | Shrubland |
| **10** | Hangklip | CAPE | 42 | 10 | 12 937 | 1.5 | 0.0 | Shrubland |
| **11** | Betty's Bay | CAPE | 21 | 17 | 17 615 | 33.1 | 0.0 | Built-up |
| **12** | Hermanus Hamilton Russel | CAPE | 22 | 7 | 11 012 | 2.3 | 18.5 | Shrubland |
| **13** | Hermanus-Onrus | CAPE | 35 | 17 | 11 152 | 1.0 | 2.5 | Shrubland |
| **14** | Hermanus-Voelklip | CAPE | 27 | 5 | 18 635 | 17.4 | 1.1 | Shrubland |
| **15** | Outeniquia Nature Reserve | GAR | 20 | 36 | 20 091 | 17.9 | 9.1 | Forest |

| | | | | | | | | |
|---|---|---|---|---|---|---|---|---|
| **16** | Denneoord | GAR | Unknown | 12 | 29 281 | 37.4 | 2.5 | Forest |
| **17** | Nelson Mandela University-North | GAR | 22 | 22 | 12 385 | 3.2 | 3.2 | Forest |
| **18** | Nelson Mandela University – Main | GAR | 13 | 31 | 12 027 | 1.9 | 10.3 | Forest |
| **19** | Victoria Bay | GAR | 40 | 6 | 13 397 | 1.4 | 26.0 | Forest |
| **20** | Wilderness Heights | GAR | 17 | 9 | 18 164 | 28.5 | 15.1 | Forest |
| **21** | Wilderness – Boosplas | GAR | Unknown | 2 | 9 192 | 0.0 | 23.9 | Forest |
| **22** | Hoekwill – Big Tree | GAR | 30 | 5 | 10 915 | 0.2 | 33.6 | Forest |
| **23** | Hoekwill – Oakhurst | GAR | 40 | 2 | 11 025 | 0.4 | 39.0 | Forest |
| **24** | Knysna – Goudveld 1 | GAR | 27 | 24 | 5 700 | 0.0 | 0.0 | Forest |
| **25** | Knysna – Goudveld 2 | GAR | 20 | 11 | 10 798 | 0.2 | 19.5 | Forest |
| **26** | Knysna – Sandberg | GAR | 100 | 34 | 9 674 | 0.1 | 54.2 | Forest |
| **27** | Sedgefield – Black Loerie | GAR | 35 | 30 | 10 660 | 0.4 | 12.9 | Forest |
| **28** | Knysna – Diepwalle | GAR | Unknown | 3 | 8 021 | 0.1 | 0.0 | Forest |
| **29** | Knysna – Pezula golf | GAR | 5 | 2 | 19 389 | 62.9 | 0.0 | Built-up |
| **30** | Plettenberg Bay | GAR | 18 | 27 | 29 120 | 62.3 | 3.6 | Built-up |
| **31** | Nature’s Valley – Farm | GAR | 30 | 14 | 9 985 | 0.3 | 16.0 | Forest |
| **32** | Nature’s Valley – Main | GAR | 11 | 21 | 10 657 | 6.1 | 6.6 | Forest |
| **33** | Nature’s Valley - Park | GAR | 22 | 19 | 9 790 | 0.1 | 0.0 | Forest |

**Supplementary Table 2.** Comparison of beta diversity and sequencing depth across replicate samples.

| Beta diversity of sequencing replicates | | | | |
|---|---|---|---|---|
| **Replicate 1** | **Replicate 2** | **Beta diversity** | **Read 1** | **Read 2** |
| CBF457a | CBF457b | **0.05** | 48 417 | 48 121 |
| CBF481a | CBF481b | **0.07** | 53 676 | 55 091 |
| CBF454a | CBF454b | **0.07** | 46 359 | 50 055 |
| CBF451a | CBF451b | **0.10** | 34 152 | 40 196 |
| CBF449a | CBF449b | **0.12** | 34 615 | 41 612 |
| CBF450b | CBF450b | **0.14** | 33 304 | 42 502 |
| CBF453a | CBF453b | **0.15** | 39 631 | 52 832 |
| CBF393a | CBF393b | **0.20** | 63 483 | 42 974 |
| CBF480a | CBF480b | **0.24** | 38 681 | 62 770 |

**Supplementary Table 3.** Results of PERMANOVA models based on weighted and unweighted UniFrac distances testing shifts in gut microbial community composition. Analyses were performed using the adonis2 function under two modelling frameworks: (A) Human Footprint and (B) land-use and dietary predictors, including built-up areas, cultivated land, $\delta^{15}N$_mean, $\delta^{15}N$_diff, $\delta^{13}C$_mean, and $\delta^{13}C$_diff. Sex and study area were included as control variables. For each explanatory variable, F-values, coefficients of determination ($R^2$), and p-values based on 9,999 permutations, restricted to between troops, are reported.

# A

| Metrics | Explanatory variables | F-value | $R^2$ | P-value |
|---|---|---|---|---|
| **Unweighted Unifrac** | Human Footprint | 8.39 | 0.018 | 0.020 |
| | Area-GAR | 18.72 | 0.047 | <0.001 |
| | Sex-Male | 3.42 | 0.007 | <0.001 |
| **Weighted Unifrac** | Human Footprint | 13.74 | 0.030 | <0.001 |
| | Area-GAR | 18.87 | 0.044 | <0.001 |
| | Sex-Male | 2.35 | 0.005 | 0.003 |

# B

| Metrics | Explanatory variables | F-value | $R^2$ | P-value |
|---|---|---|---|---|
| **Unweighted Unifrac** | Built-up | 7.65 | 0.016 | 0.016 |
| | Cultivated | 9.29 | 0.019 | 0.004 |
| | d15N_mean | 8.46 | 0.018 | 0.005 |
| | d15N_diff | 1.75 | 0.004 | 0.333 |
| | d13C_mean | 5.13 | 0.011 | 0.032 |
| | d13C_diff | 1.54 | 0.003 | 0.333 |
| | Area-GAR | 18.72 | 0.033 | <0.001 |
| | Sex-Male | 3.42 | 0.033 | <0.001 |
| **Weighted Unifrac** | Built-up | 10.07 | 0.021 | 0.017 |
| | Cultivated | 8.34 | 0.017 | 0.035 |
| | d15N_mean | 7.81 | 0.016 | 0.053 |
| | d15N_diff | 1.30 | 0.003 | 0.143 |
| | d13C_mean | 4.54 | 0.009 | 0.460 |
| | d13C_diff | 1.33 | 0.003 | 0.131 |
| | Area-GAR | 15.77 | 0.033 | <0.001 |
| | Sex-Male | 2.26 | 0.005 | 0.003 |

**Supplementary Table 4.** Results of dbRDA models based on weighted and unweighted UniFrac distances, testing shifts in gut microbial community composition while accounting for spatial structure. Analyses were performed using data averaged at the troop level to account for non-independence among samples. Two modelling frameworks were considered: (A) Human Footprint and (B) land-use and dietary predictors, including built-up areas, cultivated land, $\delta^{15}N$_mean, $\delta^{15}N$_diff, $\delta^{13}C$_mean, and $\delta^{13}C$_diff. Sex and study area were included as control variables, and dbMEMs were added to account for spatial structure. For each explanatory variable, F-values and p-values based on 9,999 permutations are reported.

# A

| Metrics | Explanatory variables | F-value | P-value |
|---|---|---|---|
| Unweighted Unifrac | Human Footprint | 1.26 | 0.162 |
| | Area-GAR | 0.67 | 0.795 |
| | Sex-Male | 1.19 | 0.208 |
| Weighted Unifrac | Human Footprint | 2.17 | 0.011 |
| | Area-GAR | 0.88 | 0.507 |
| | Sex-Male | 1.45 | 0.098 |

# B

| Metrics | Explanatory variables | F-value | P-value |
|---|---|---|---|
| Unweighted Unifrac | Built-up | 1.08 | 0.319 |
| | Cultivated | 1.15 | 0.269 |
| | d15N | 1.58 | 0.039 |
| | d13C | 1.85 | 0.015 |
| | Area-GAR | 0.71 | 0.738 |
| | Sex-Male | 1.27 | 0.146 |
| Weighted Unifrac | Built-up | 1.59 | 0.071 |
| | Cultivated | 1.08 | 0.331 |
| | d15N | 1.56 | 0.072 |
| | d13C | 0.92 | 0.516 |
| | Area-GAR | 1.07 | 0.334 |
| | Sex-Male | 1.39 | 0.123 |

**Supplementary Table 5.** Linear mixed model results testing variation in gut microbial community dispersion among chacma baboon troops. Distances to group centroids were calculated using PERMDISP2 test (*betadisper)* based on weighted and unweighted UniFrac and then used as response variables in the models. Models included Human Footprint, sex, and study area as covariates. Reported estimates and 95% confidence intervals correspond to model-averaged coefficients across models (ΔAICc < 2). Analyses were restricted to troops with >10 samples (21 troops, n = 379 samples).

| Metrics | Predictors | Estimates | 95% confidence intervals |
|---|---|---|---|
| **Unweighted Unifrac** | Human Footprint | 0.004 | [-0.008; 0.015] |
| | Area-GAR | 0.051 | [0.029; 0.074] |
| | Sex-Male | 0.027 | [0.014; 0.039] |
| **Weighted Unifrac** | Human Footprint | 0.000 | [-0.001; 0.001] |
| | Area-GAR | -0.001 | [-0.003 ; 0.001] |
| | Sex-Male | -0.001 | [- 0.003; 0.001] |

**Supplementary Table 6.** ANCOM-BC results of taxonomic and functional differential abundance in chacma baboon along different anthropogenic variables. Taxonomic differential abundances associated with Human Footprint Index (HFI) were presented at the (A) phylum, (B) family, (C) genus, and (D) ASV levels. Taxonomic differential abundances associated with land-use variables (built-up and cultivated areas) and stable isotopes at the between- and within-troop levels ($\delta^{15}$N_mean/ $\delta^{13}$C_mean and $\delta^{15}$N_diff/$\delta^{13}$C_diff, respectively) were included at the (E) phylum, (F) family, (G) genera, and (H) ASV levels. MetaCyc pathway differential abundances were included for the (I) HFI model, and (J) land-use and isotope models. (K) Enzyme commission (EC) differential abundances for HFI model. All models included sex and study area as covariates. Only significant features are reported at family, genus, ASV, pathway, and EC levels, while all phyla are included. Tables report log fold change (LFC), standard error (SE), test statistic (W), p-values, adjusted q-values, and significance before (diff) and after sensitivity filtering (diff_robust). P-values were corrected using the Benjamin–Hochberg procedure, and taxa with adjusted q-values < 0.05 were considered differentially abundant.

Files provided separately as Supplementary_Table_ 6.zip.

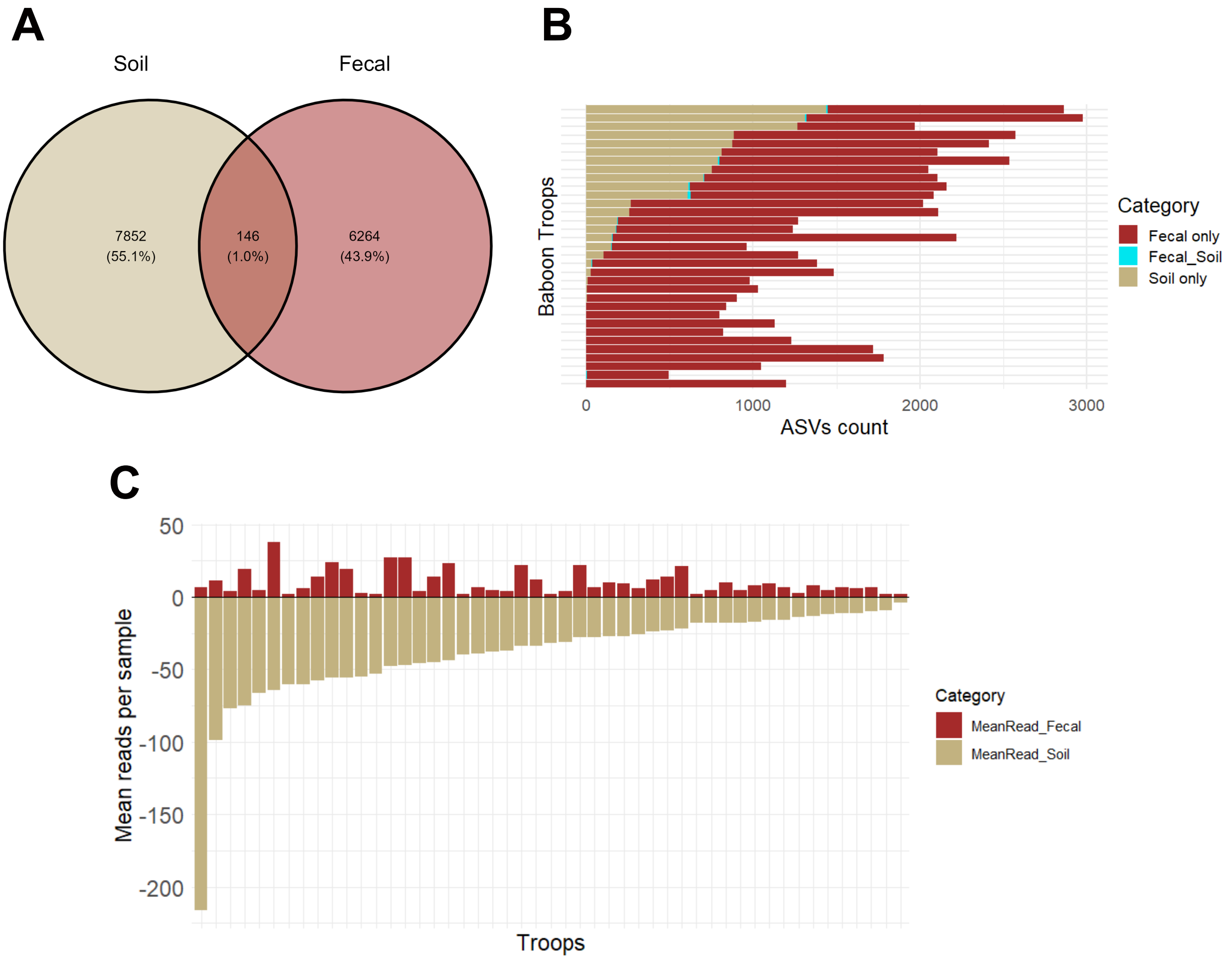


**Supplementary Figure 1.** Comparison of ASV composition between soil and fecal samples and assessment of potential environmental contamination. A. Venn diagram showing the number and proportion of ASVs detected in fecal and soil samples. A total of 146 ASVs were shared between sample types (1.0%), compared to 7,852 soil-specific ASVs (55.1%) and 6,264 fecal-specific ASVs (43.9%). B. Distribution of ASVs across baboon troops, showing the number of ASVs detected exclusively in fecal samples (red), exclusively in soil samples (beige), or shared between both sample types (blue). At the troop level, only 0–15 ASVs were shared between soil and fecal samples. C. For shared ASVs, comparison of mean read counts between fecal and soil samples. ASVs more abundant in soil than in feces were identified as potential environmental contaminants. Based on these criteria, 57 shared ASVs that were more abundant in soil than in feces were removed from downstream analyses. Most of these ASVs were of low abundance and would have been excluded by subsequent filtering steps.

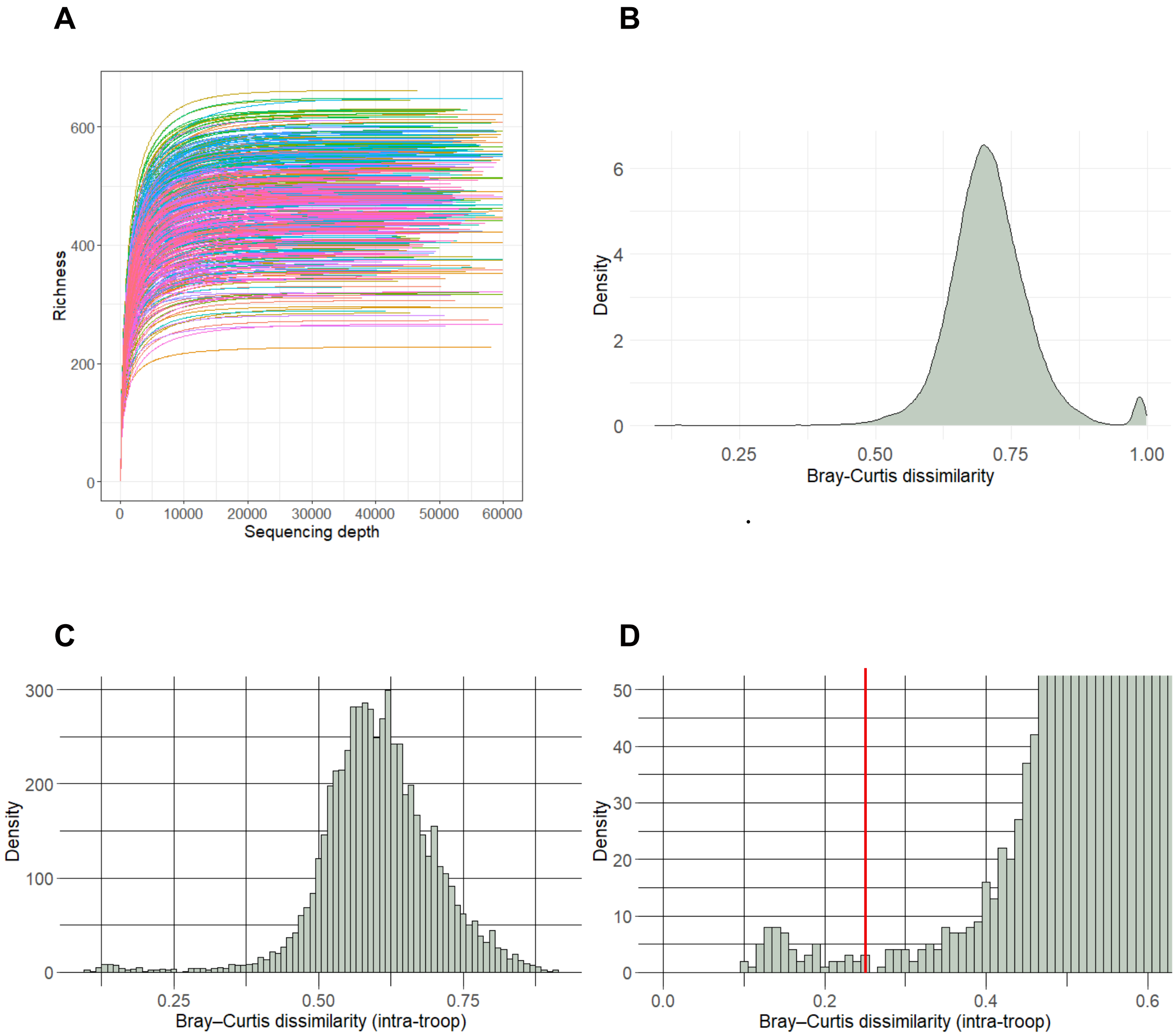


**Supplementary Figure 2. Rarefaction curves and beta-diversity structure used to define sequencing depth threshold and identify duplicates.** A. Rarefaction curves for all fecal samples (N = 504). Each line represents a sample, and the red vertical line indicates the rarefaction depth selected (25 000 reads), based on the plateauing of richness across samples. B. Distribution of pairwise Bray–Curtis dissimilarities across all samples. The main peak corresponds to inter-individual variation in gut microbial communities, whereas the smaller peak represents highly dissimilar comparisons involving non-baboon samples. C. Density distribution of Bray–Curtis dissimilarities across intra-troop sample pairs, showing a clear separation between duplicate samples (low dissimilarity) and distinct individuals (higher dissimilarity). D. Zoom on the low-dissimilarity range, highlighting a secondary peak corresponding to highly similar samples. The red vertical line (0.25) indicates the threshold used to identify duplicate samples from the same individual.

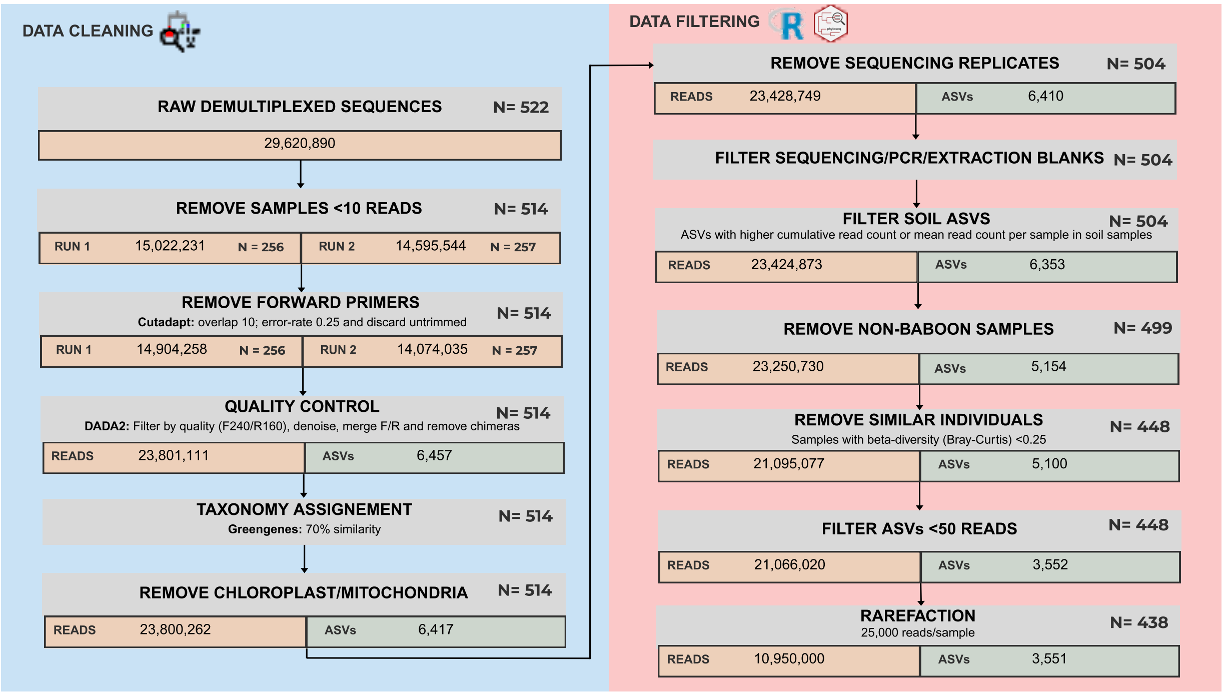


**Supplementary Figure 3.** Bioinformatic filtering workflow and sample/sequence retention across processing steps. Overview of the bioinformatic pipeline applied to fecal samples, from raw demultiplexed sequences to the final dataset used for downstream analyses. The workflow is divided into two main stages: data cleaning (left panel) and data filtering (right panel). During data cleaning, raw sequences (N = 522 samples) were processed through read filtering, primer trimming, denoising (DADA2), taxonomic assignment, and removal of chloroplast and mitochondrial sequences, resulting in 514 samples and 6 417 ASVs. Subsequent filtering steps included the removal of low-depth samples (<10 reads), sequencing and extraction blanks, ASVs associated with soil contamination, non-baboon samples, and duplicate samples identified based on Bray–Curtis dissimilarity, followed by removal of low-abundance ASVs (<50 reads). Finally, samples were rarefied to 25,000 reads per sample, yielding a final dataset of 438 fecal samples and 3 551 ASVs. At each step, the numbers of retained samples (N), reads, and ASVs are indicated.

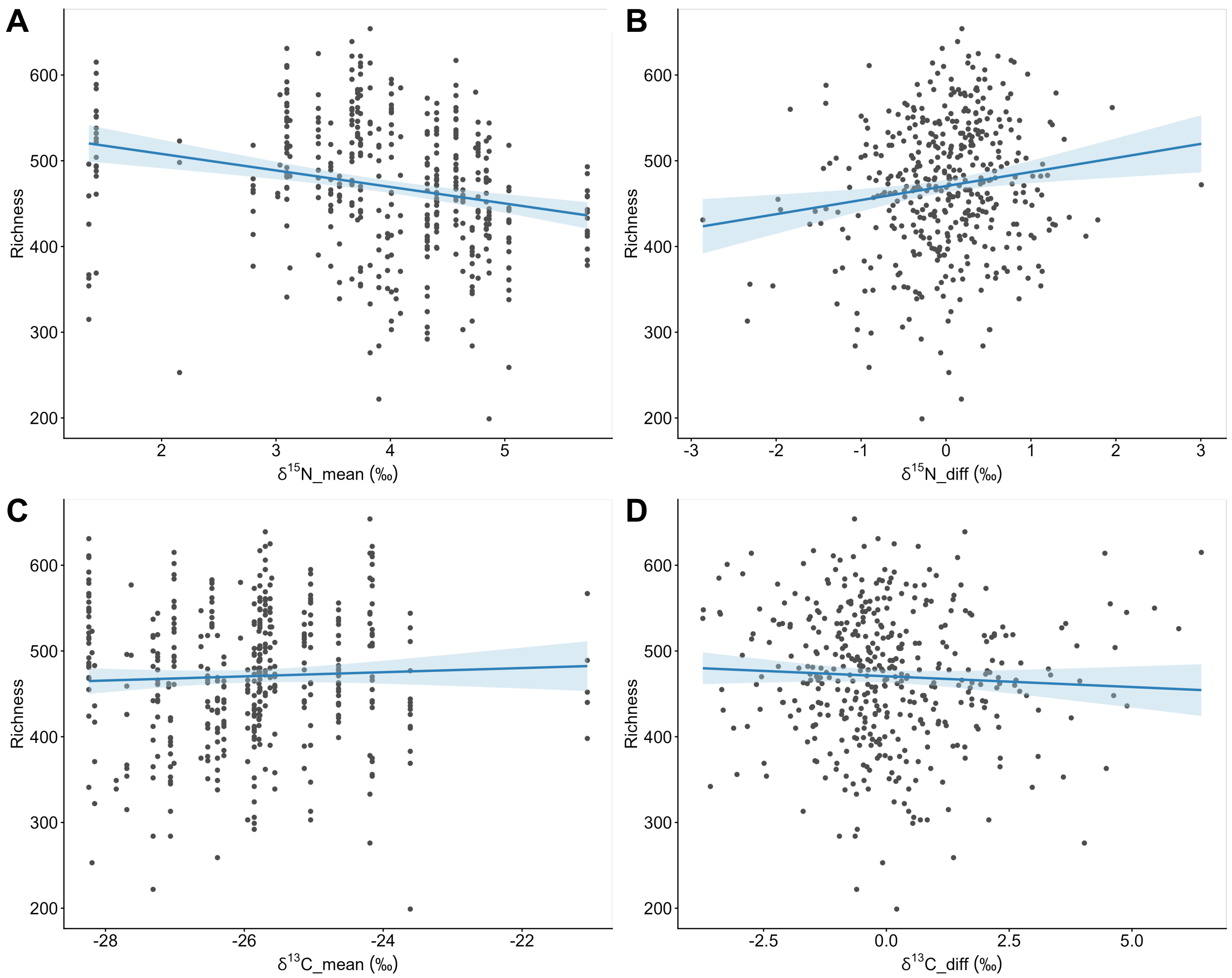


**Supplementary Figure 4.** Relationships between stable isotope values and gut microbial alpha diversity, illustrating the decomposition of $\delta^{15}N$ effects. A. Relationship between $\delta^{15}N$_mean (between-troop variation) and richness, showing a negative association across troops. B. Relationship between $\delta^{15}N$_diff (within-troop variation) and richness, showing a positive association within troops. C. Relationship between $\delta^{13}C$_mean and richness, showing no strong association at the between-troop level. D. Relationship between $\delta^{13}C$_diff and richness, showing no consistent within-troop effect.

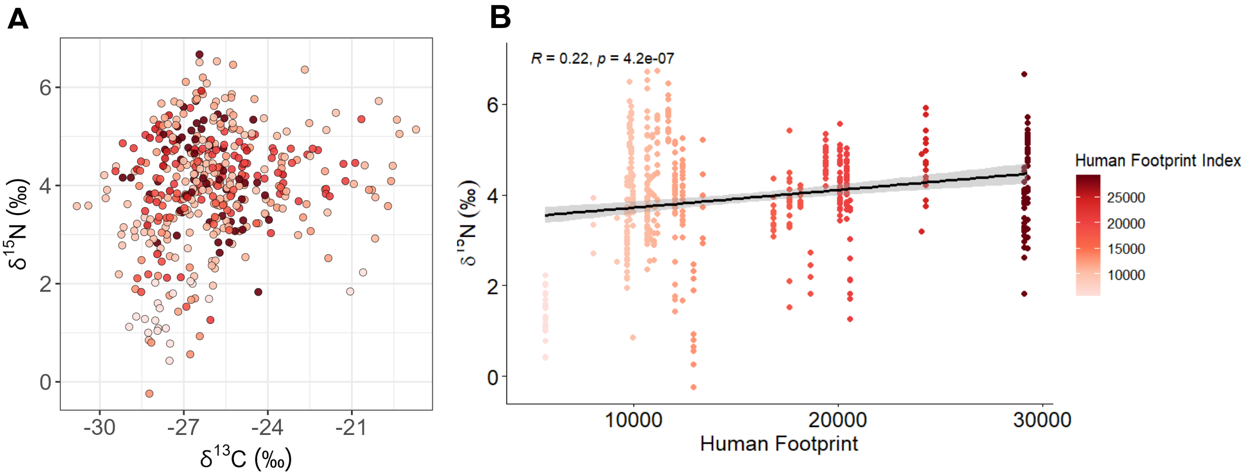


**Supplementary Figure 5.** Distribution of stable isotope values ($\delta^{13}C$ and $\delta^{15}N$) across samples along the anthropogenic gradient. A. Scatterplot showing the relationship between $\delta^{13}C$ and $\delta^{15}N$ values across all fecal samples. B. Relationship between $\delta^{15}N$ and Huma Footprint Index, showing a positive association. Points are coloured according to the Human Footprint Index (HFI), with darker colours indicating higher levels of anthropogenic pressure. Fitted values from linear model is shown as solid line, with shaded areas representing 95% confidence intervals.

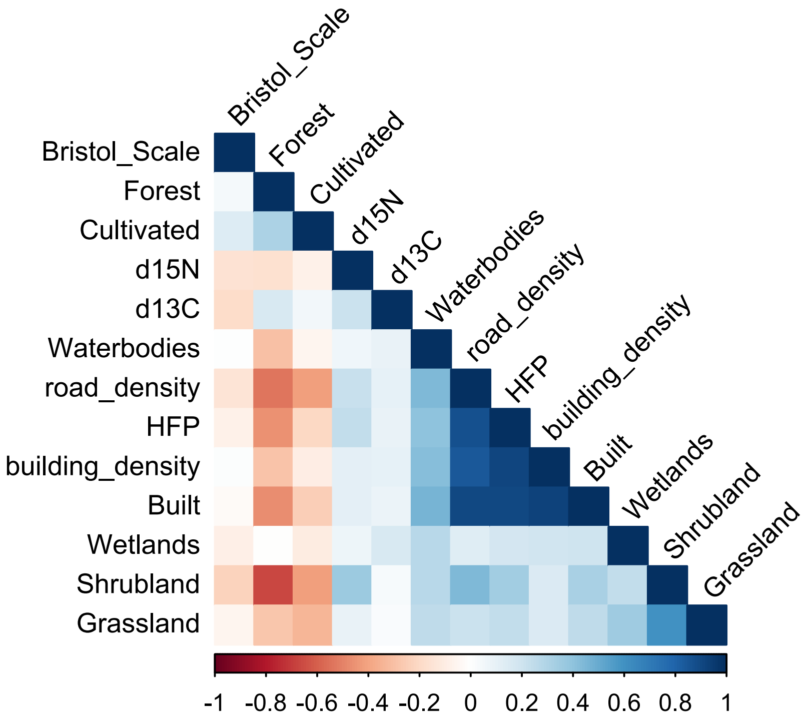


**Supplementary Figure 6.** Pairwise Spearmann correlations among variables considered prior the analysis.

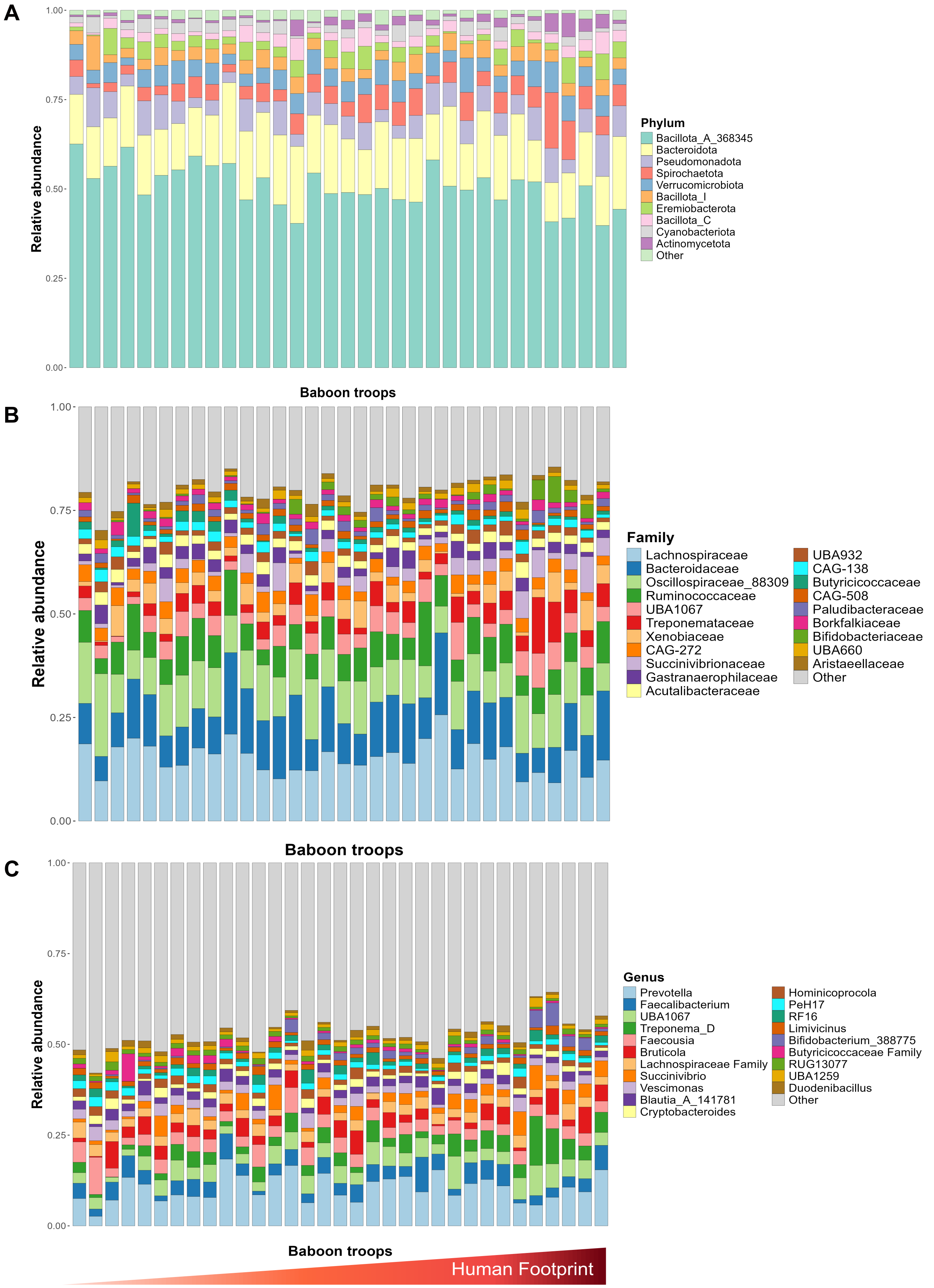


**Supplementary Figure 7.** Taxonomic composition of the gut microbiome across baboon troops along the anthropogenic gradient. Relative abundance of dominant taxa at the (A) phylum, (B) family, and (C) genus levels across baboon troops. Each bar represents one troop, and taxa are shown as relative proportions of total reads. Baboon troops are ordered along the Human Footprint Index (HFI) gradient (from low to high anthropogenic pressure, left to right). Only the most abundant taxa are displayed, with remaining taxa grouped as “Other”.

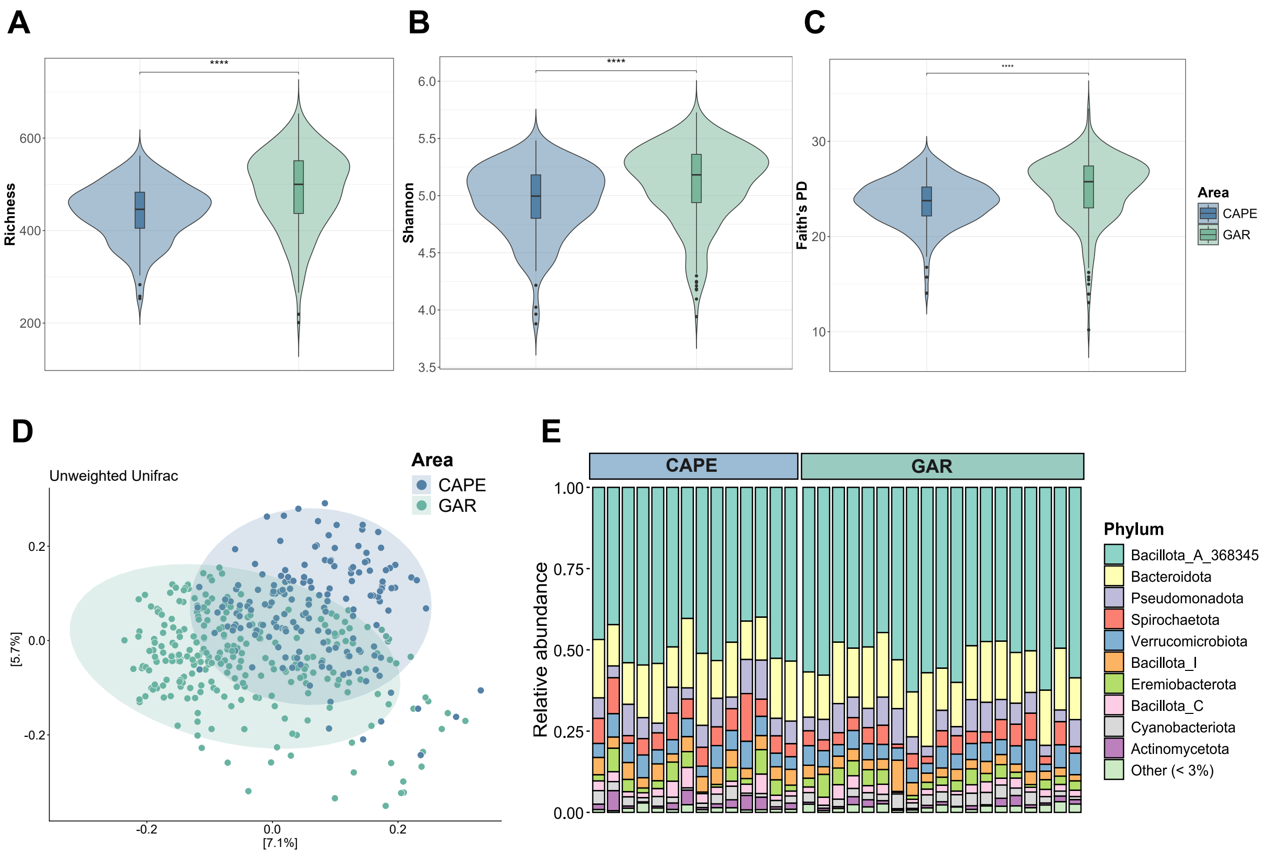


**Supplementary Figure 8:** Differences between areas (CAPE/GAR) in alpha, beta diversity and microbial differential abundance. Variation in (A) observed richness, (B) Shannon diversity, and (C) Faith's phylogenetic diversity between CAPE and GAR areas. **** indicates significance difference ($p<0.01$) based on Wilcoxon rank-sum test. (D) Beta diversity pattern is represented using PCoA based on unweighted UniFrac. Colours corresponds to baboon location of the samples. (E) Mean relative abundances of dominant taxa across baboon troops at the phylum level.

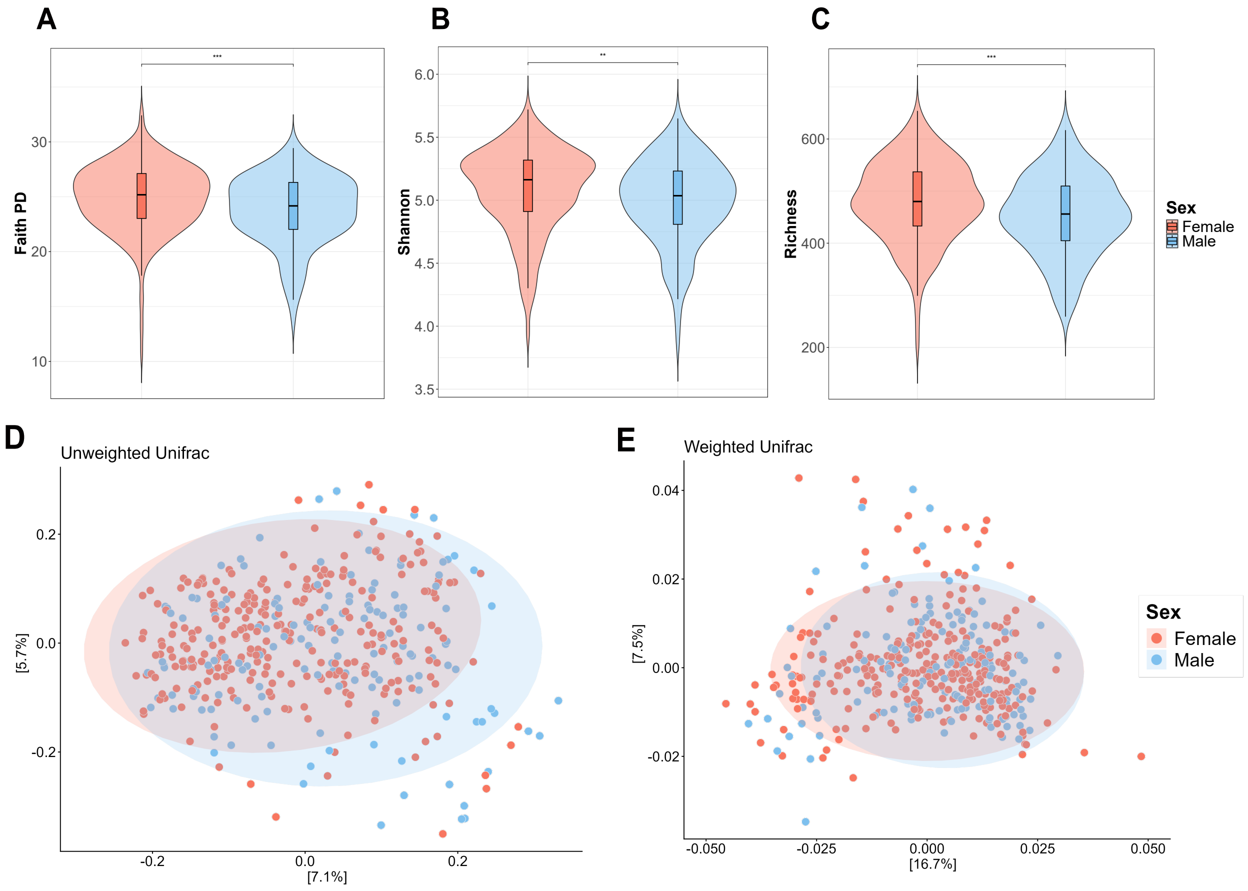


**Supplementary Figure 9:** Sex differences in alpha, beta diversity and microbial differential abundance in baboons. Variations in (A) observed richness, (B) Shannon diversity, and (C) Faith's phylogenetic diversity between male and female. ** indicates significance difference (p<0.05) and *** (p<0.01) based on Wilcoxon rank-sum test. (D) Beta diversity patterns are represented using PCoA based on (D) unweighted UniFrac, and (E) weighted UniFrac. Colours corresponds to baboon sex.

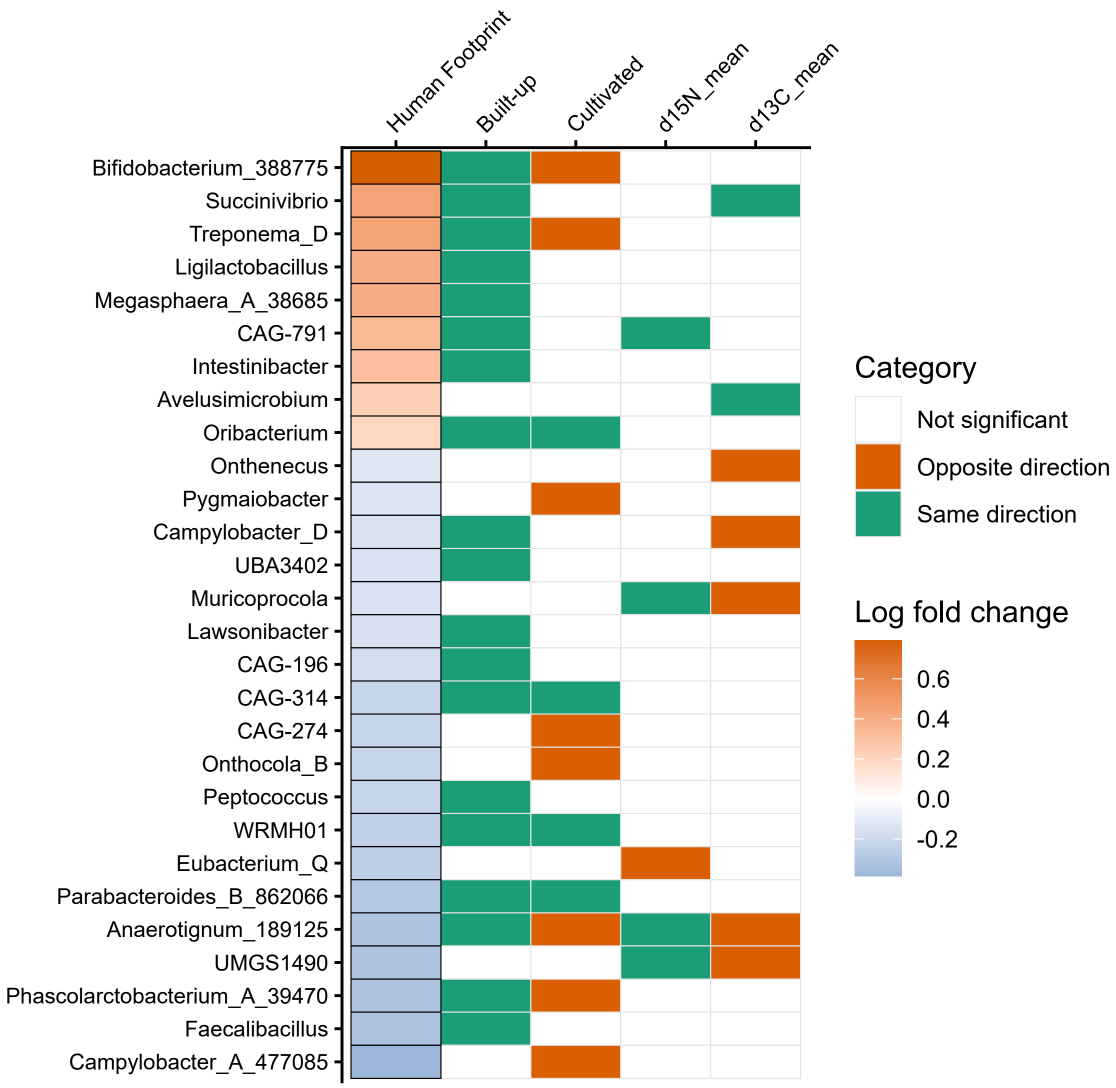


**Supplementary Figure 10:** Shared microbial genera showing significant differential abundance between Human Footprint (HFI) and other anthropogenic variables. The HFI column shows Log Fold Change (LFC) magnitude (red: increase; blue: decrease). Subsequent columns indicate response direction relative to HFP with green for same direction (both increasing or both decreasing), red for opposite direction, and white for non-significant differential abundance ($p > 0.05$). Only genera significantly associated with the Human Footprint Index (HFI) and at least one other predictor are represented.
